# Predicting Experimental Success in De Novo Binder Design: A Meta-Analysis of 3,766 Experimentally Characterised Binders

**DOI:** 10.1101/2025.08.14.670059

**Authors:** Max D. Overath, Andreas S. H. Rygaard, Christian P. Jacobsen, Valentas Brasas, Oliver Morell, Pietro Sormanni, Timothy P. Jenkins

## Abstract

Designing high-affinity de novo protein binders has become increasingly tractable, yet *in vitro* prioritisation continues to depend on heuristics in the absence of systematic analysis. Here, we present a large-scale meta-analysis of 3,766 experimentally tested binders across 15 structurally diverse targets. Using a unified, high-throughput pipeline that re-predicts each binder-target complex with AF2 (initial guess and ColabFold), AF3 and Boltz-1, we extract over 200 structural, energetic and confidence features per design. We show that interface-focused metrics, most notably the AF3-derived interaction prediction Score from Aligned Errors (ipSAE) outperform commonly used scores such as ipAE and ipTM, with a significant 1.4-fold increase in average precision compared to ipAE. We further show that combining these metrics with orthogonal physicochemical interface descriptors, including Rosetta ΔG/ΔSASA and interface shape complementarity, improves predictive performance. While overall performance varies by target, simple linear models trained on a small number of AF3-derived features generalize well across datasets. We propose interpretable, target-agnostic filtering strategies, such as combining AF3 ipSAE_min rankings with structural filters, to improve precision in selecting binders for testing. Finally, we release the complete dataset establishing a community resource to benchmark and accelerate de novo binder discovery.

## Introduction

Recent advances in de novo protein binder design have opened new possibilities for computational protein engineering. Methods such as RFdiffusion (1), BindCraft (2), and AlphaProteo (3) now enable the generation of high-affinity protein binders directly from target structures, without relying on natural templates. These methods have already been applied across a wide spectrum of use cases (4–13), highlighting their potential in therapeutics, diagnostics, and basic research.

The synthesis and screening of large libraries of protein binders remain costly and labour intensive. Consequently, the *in silico* filtering of candidate designs has become a central challenge. Filtering de novo designed proteins is especially challenging, as it usually requires selection among a highly similar pool of structures.

A major advance in de novo binder design was the discovery that deep learning–based protein structure prediction models especially AlphaFold2 (AF2) (14) can effectively prioritize candidates prior to experimental validation, increasing the experimental success rate. Confidence metrics derived from AF2 predictions, such as per-residue confidence (pLDDT), interaction predicted aligned error (ipAE), and interface predicted TM-score (ipTM), have been shown to be predictive of *in vitro* binding, outperforming traditional physics-based scores such as Rosetta energies (15).

Despite this progress, design success remains very inconsistent, and there are no standard criteria to prioritise binders for experimental testing. Since the adoption of AF2-based filtering, several newer and more accurate protein structure prediction models have emerged, including AlphaFold3 (AF3) (16), Boltz-1 (17), Boltz-2 (18) and Chai-1 (19). However, their performance in prediction of *in vitro* binding has not been systematically evaluated. In parallel, a growing number of alternative metrics and filters have been proposed to improve scoring of protein–protein interactions (10, 20–26), yet their generalisability across large, diverse datasets remains unclear. This is corroborated by most de novo campaigns producing only a small number of validated binders and typically focusing on related targets and thus limiting the ability to benchmark scoring methods at scale.

To tackle these limitations, we curated a dataset of over 3,700 experimentally tested de novo binders targeting 15 structurally and functionally diverse proteins. To score these designs efficiently at scale, we developed a streamlined pipeline that re-predicts each binder–target complex using four structure prediction tools: the initial guess and ColabFold implementations of AF2 (15, 27), as well as AF3 and Boltz-1. We benchmarked the impact of multiple sequence alignment (MSA) generation, number of recycles, and number of models on both runtime and prediction accuracy, identifying configurations that substantially improve throughput with minimal loss in model confidence. Together, we extracted over 200 structural, energetic, confidence, and sequence-based features per design.This large-scale meta-analysis enabled us to systematically evaluate which metrics best predict experimental success across targets, design protocols, and prediction methods.

## Results

### Overview of data

To assemble a dataset spanning diverse target classes and binder designs, we curated experimentally tested de novo binders from multiple published sources (Table 1). In total, we collected data from binders covering 15 distinct targets, including receptor tyrosine kinases (RTKs), pathogen-derived antigens, immune modulators, and a range of additional targets such as a snake venom short-chain *α*-neurotoxin (sntx), a proto-oncogene, and two peptide MHC (pMHC) complexes. Mapping the target interaction residues for true experimental binders shows that their binding sites are highly conserved (Fig. **1**A).

**Figure 1.**
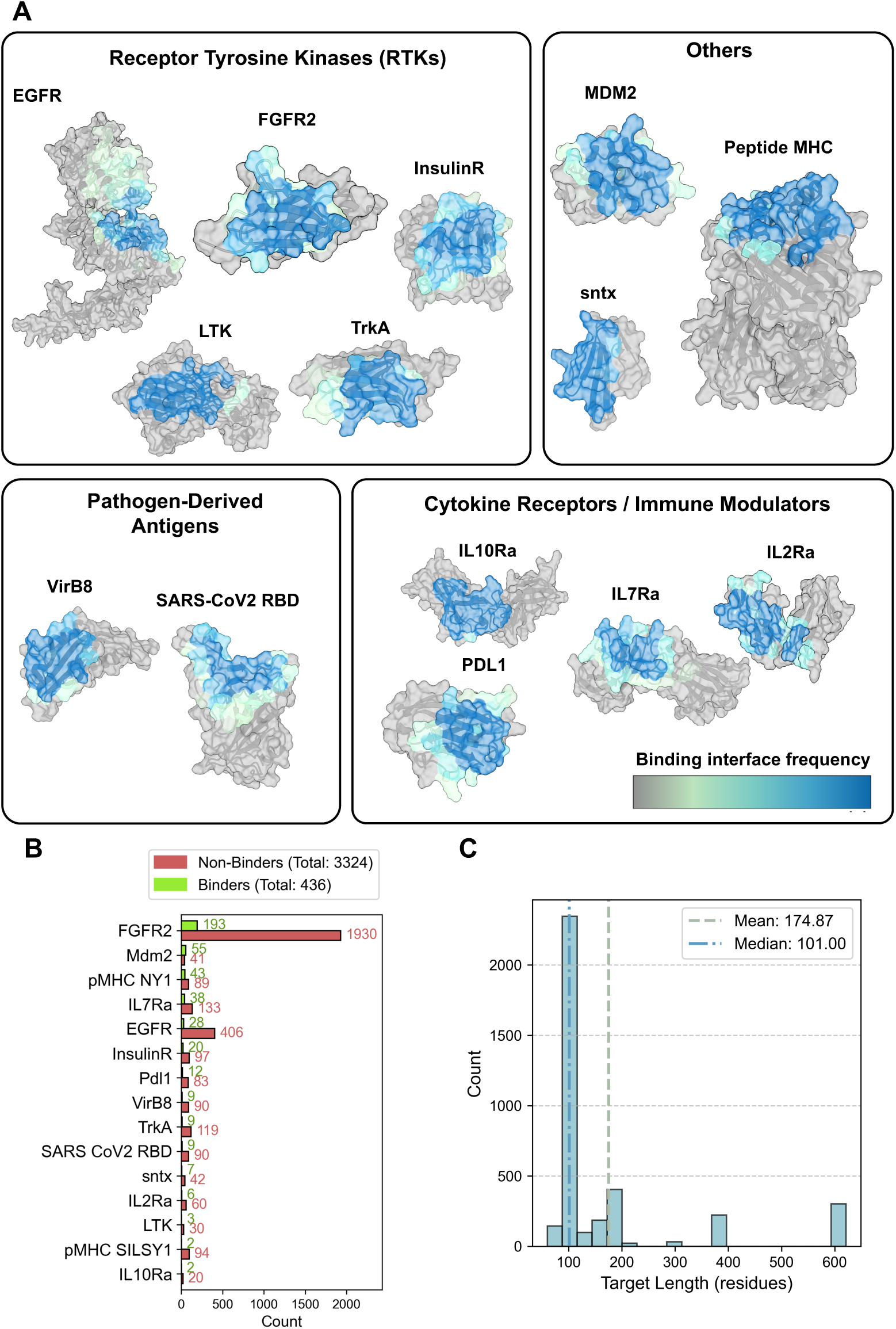
Overview of binder data: **A)** A broad range of targets were selected, for which binder-target complex structures linked to *in vitro* data were available. These targets include receptor tyrosine kinases (RTKs), pathogen-derived antigens, immune modulators, and others including a snake venom short-chain *α*-neurotoxin (sntx), a proto-oncogene and two peptide major histocompatibility complexes (pMHCs) (only one pMHC structure shown for reference). The coloring indicates the frequency of a given residue being part of the binding interface for all true binders. **B)** Count of binders and non-binders for each target in the dataset. Number of binders differs significantly across targets both in absolute and relative terms. **C)** Histogram of target lengths in compiled dataset.

**Table 1:**
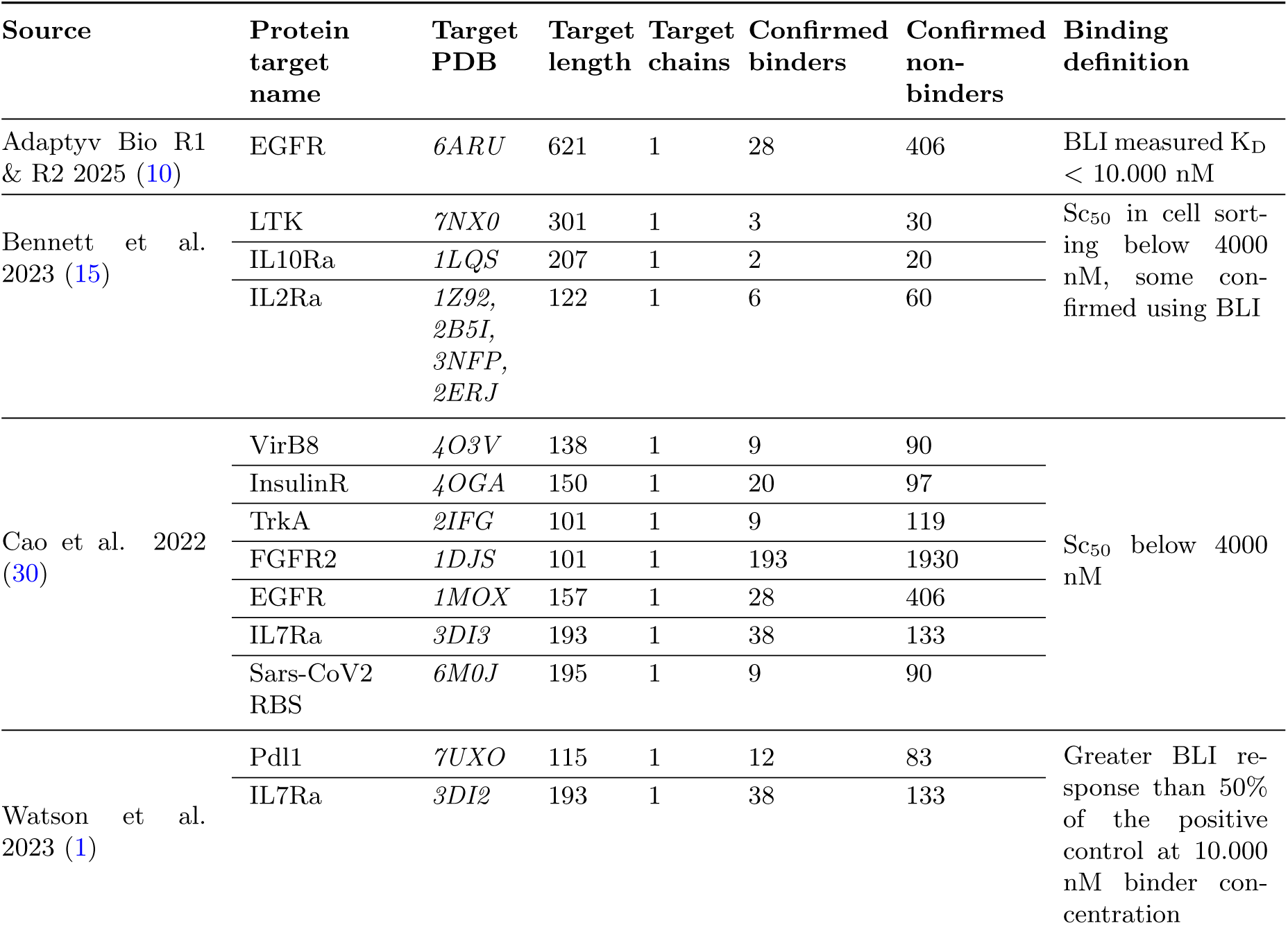
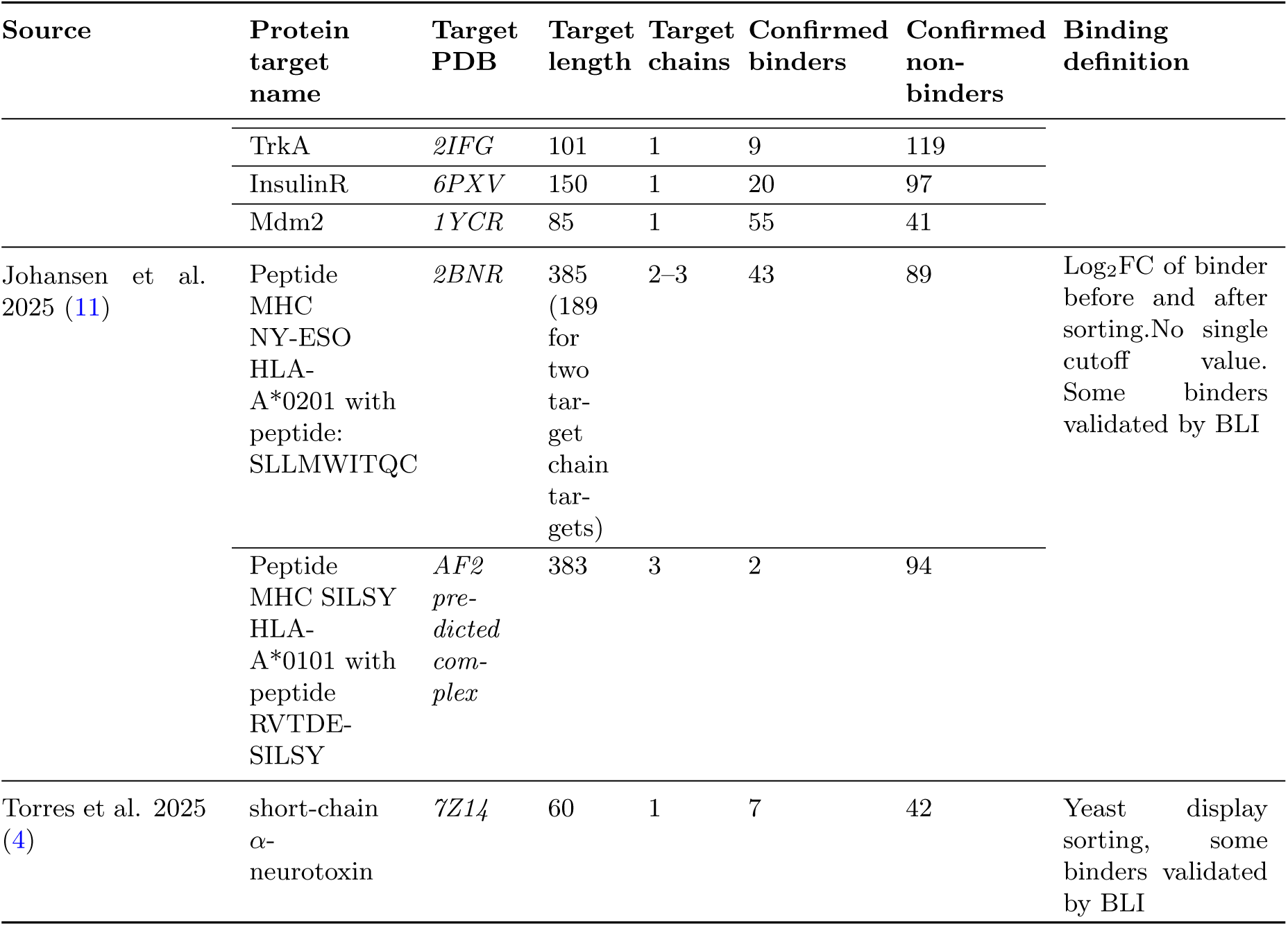
List of protein targets, their PDB identifiers, numbers of confirmed binders and non-binders, and binding definitions.

**Table 2:**
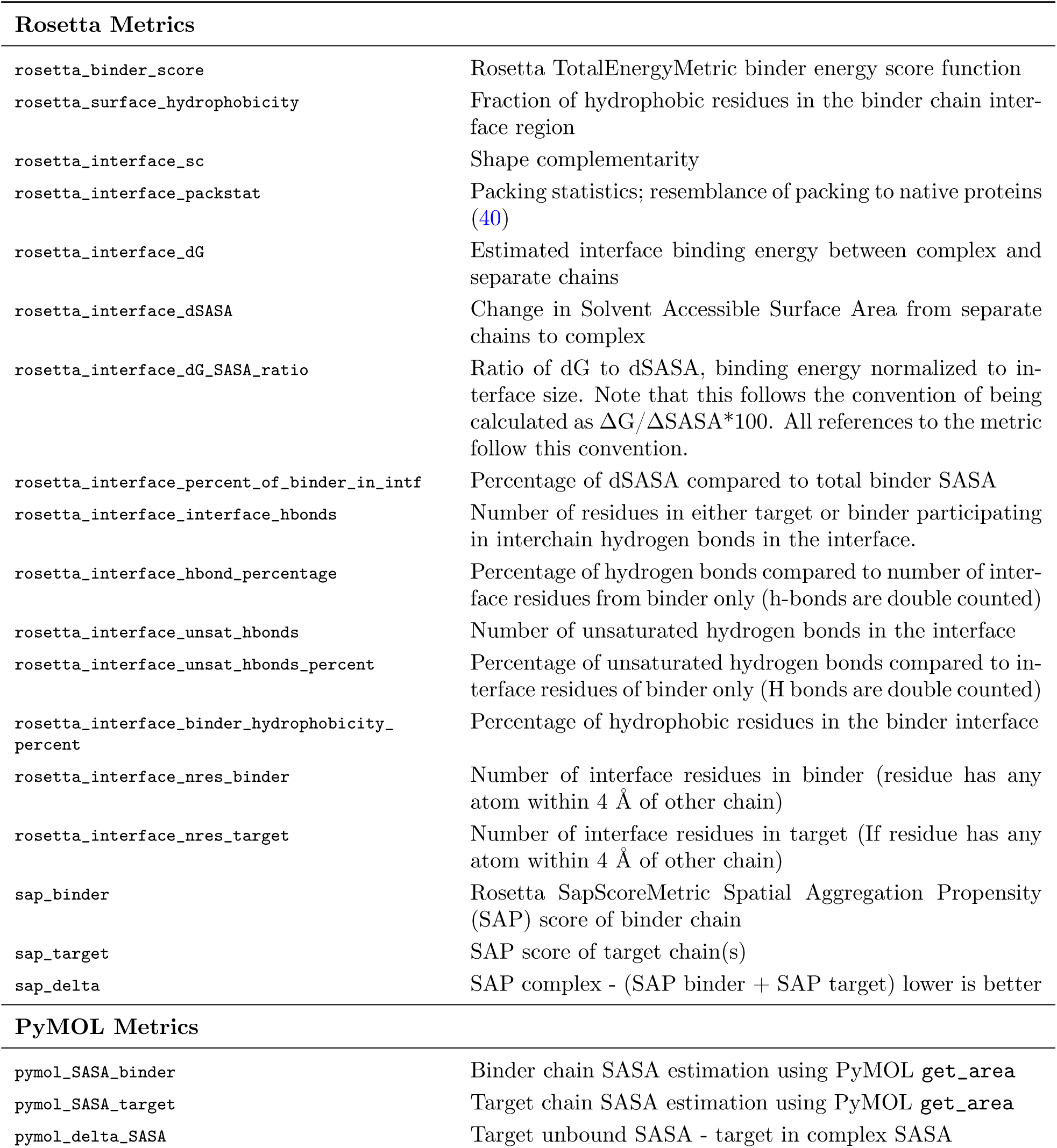
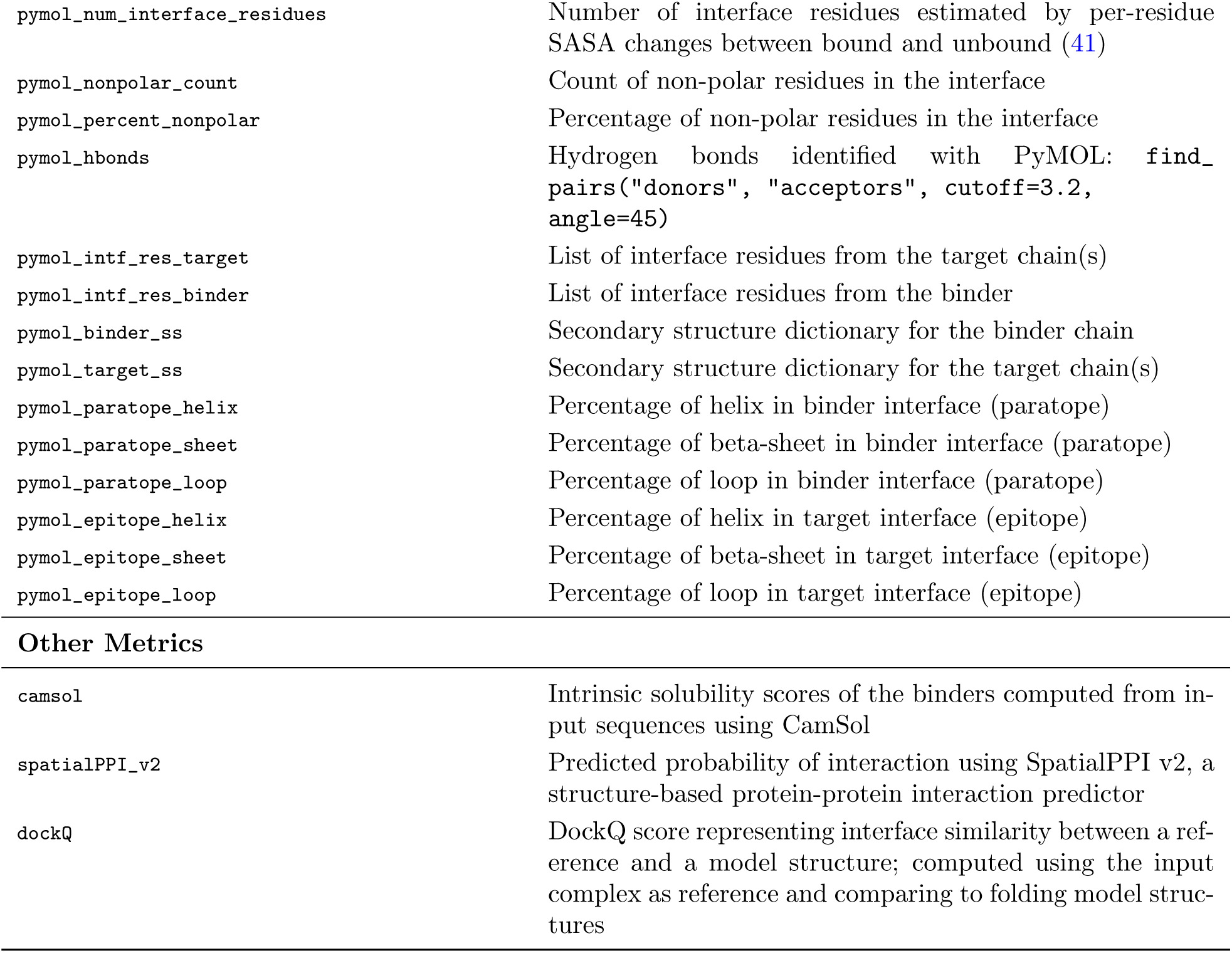
Overview of computed interface metrics and corresponding description

| Rosetta Metrics |  |
| --- | --- |
| rosetta_binder_score | Rosetta TotalEnergyMetric binder energy score function |
| rosetta_surface_hydrophobicity | Fraction of hydrophobic residues in the binder chain interface region |
| rosetta_interface_sc | Shape complementarity |
| rosetta_interface_packstat | Packing statistics; resemblance of packing to native proteins (40) |
| rosetta_interface_dG | Estimated interface binding energy between complex and separate chains |
| rosetta_interface_dSASA | Change in Solvent Accessible Surface Area from separate chains to complex |
| rosetta_interface_dG_SASA_ratio | Ratio of dG to dSASA, binding energy normalized to interface size. Note that this follows the convention of being calculated as $\Delta G / \Delta SASA * 100$. All references to the metric follow this convention. |
| rosetta_interface_percent_of_binder_in_intf | Percentage of dSASA compared to total binder SASA |
| rosetta_interface_interface_hbonds | Number of residues in either target or binder participating in interchain hydrogen bonds in the interface. |
| rosetta_interface_hbond_percentage | Percentage of hydrogen bonds compared to number of interface residues from binder only (h-bonds are double counted) |
| rosetta_interface_unsat_hbonds | Number of unsaturated hydrogen bonds in the interface |
| rosetta_interface_unsat_hbonds_percent | Percentage of unsaturated hydrogen bonds compared to interface residues of binder only (H bonds are double counted) |
| rosetta_interface_binder_hydrophobicity_percent | Percentage of hydrophobic residues in the binder interface |
| rosetta_interface_nres_binder | Number of interface residues in binder (residue has any atom within 4 Å of other chain) |
| rosetta_interface_nres_target | Number of interface residues in target (If residue has any atom within 4 Å of other chain) |
| sap_binder | Rosetta SapScoreMetric Spatial Aggregation Propensity (SAP) score of binder chain |
| sap_target | SAP score of target chain(s) |
| sap_delta | SAP complex - (SAP binder + SAP target) lower is better |
| PyMOL Metrics |  |
| pymol_SASA_binder | Binder chain SASA estimation using PyMOL `get_area` |
| pymol_SASA_target | Target chain SASA estimation using PyMOL `get_area` |
| pymol_delta_SASA | Target unbound SASA - target in complex SASA |
| `pymol_num_interface_residues` | Number of interface residues estimated by per-residue SASA changes between bound and unbound (41) |
| `pymol_nonpolar_count` | Count of non-polar residues in the interface |
| `pymol_percent_nonpolar` | Percentage of non-polar residues in the interface |
| `pymol_hbonds` | Hydrogen bonds identified with PyMOL: `find_pairs("donors", "acceptors", cutoff=3.2, angle=45)` |
| `pymol_intf_res_target` | List of interface residues from the target chain(s) |
| `pymol_intf_res_binder` | List of interface residues from the binder |
| `pymol_binder_ss` | Secondary structure dictionary for the binder chain |
| `pymol_target_ss` | Secondary structure dictionary for the target chain(s) |
| `pymol_paratope_helix` | Percentage of helix in binder interface (paratope) |
| `pymol_paratope_sheet` | Percentage of beta-sheet in binder interface (paratope) |
| `pymol_paratope_loop` | Percentage of loop in binder interface (paratope) |
| `pymol_epitope_helix` | Percentage of helix in target interface (epitope) |
| `pymol_epitope_sheet` | Percentage of beta-sheet in target interface (epitope) |
| `pymol_epitope_loop` | Percentage of loop in target interface (epitope) |
| Other Metrics |  |
| `camsol` | Intrinsic solubility scores of the binders computed from input sequences using CamSol |
| `spatialPPI_v2` | Predicted probability of interaction using SpatialPPI v2, a structure-based protein-protein interaction predictor |
| `dockQ` | DockQ score representing interface similarity between a reference and a model structure; computed using the input complex as reference and comparing to folding model structures |

Across all datasets, we identified 3,766 unique binder designs with associated experimental binding data, which had low sequence similarity within and across datasets (Fig. S1 B). Of these, 436 (11.6%) were classified as *in vitro* binders. The number of tested designs and proportion of confirmed binders varied substantially between targets, resulting in a highly imbalanced dataset (Fig. **1**B). Notably, binding definitions were not standardised across studies, with variation in affinity thresholds and assay formats (Table 1). Furthermore, the target proteins ranged from 60 to 621 residues in length (mean: 174; median: 101), reflecting the broad target diversity of the dataset (Fig. **1**C).

### Pipeline setup for time efficient scoring of binder complexes

To enable high-throughput evaluation of binder designs, we developed a scalable and automated pipeline that integrates four structure prediction tools and extracts over 200 structural, sequence-based, and energetic features per design from both the predicted complexes and input structures (Fig. **2**A). A major bottleneck in de novo binder design is the limited throughput of structure prediction tools used to evaluate candidate designs. To test the impact of key runtime parameters, we curated a benchmark dataset consisting of 56 experimentally validated true binders sampled across all 15 targets. Twelve of these binders had experimentally resolved structures in complex with the target, which were used to evaluate accuracy of structure predictors across different settings. The pipeline was evaluated under three configurations (maximum, intermediate, and minimal) that differed in the computational rigor of their settings, such as the extent of multiple sequence alignment (MSA) generation, number of recycles, and number of model runs per prediction for ColabFold, Boltz-1 and AF3 (Fig. **2**B). The AF2 initial guess implementation was excluded from this optimisation, as it was already designed for high-throughput scoring and served as a reference in this benchmarking.

**Figure 2.**
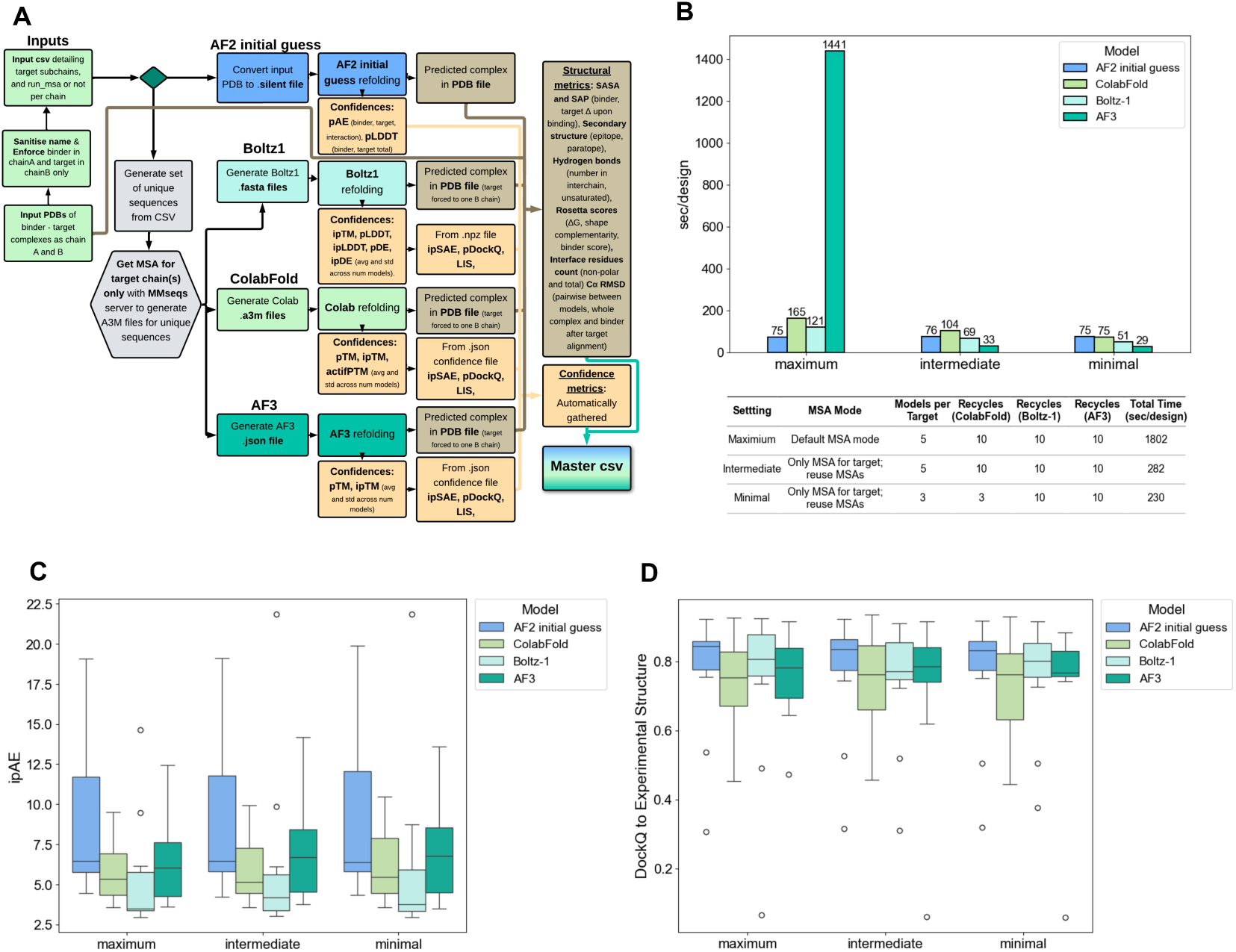
Scoring pipeline overview and set-up: **A)** The scoring pipeline takes binder-target complex PDBs as inputs with the binder as chain A and the target as chain B. These are paired with an input CSV, which specifies the subchains of the target and multiple sequence alignemnt (MSA) settings. For the AF2 initial guess, the input PDB is used to fix the target structure, while only the binder is re-predicted. For ColabFold, Boltz-1, and AF3, the binder sequence is extracted together with the relevant target subchains, and the entire complex is re-predicted. For this, a MSA is generated once for each unique target chain across the entire input set using MMseqs2. The MSA files are then written into the specific input format required by each folding model. From each model, confidence scores are extracted from output files, and ipSAE, pDockQ, pDockQ2, ipAE and LIS are calculated for all except AF2 initial guess. Additionally, the highest-confidence predicted complex for each model is extracted and converted to a PDB file if needed. These files are collected and a range of structural metrics are calculated for each complex in addition to pairwise RMSDs between the models. **B)** Time comparison of structure prediction models using different settings. Using the minimal settings significantly decreases overall time per complex prediction. Table shows different settings that were tested per structure prediction model. **C)** ipAE of models across different pipeline settings show no significant decrease in model confidences. **D)** DockQ of predicted complexes to experimental structures show no significant decrease in alignment to experimental structures across models.

The average time required to run all model predictions per design decreased substantially, from 1802 seconds using the maximum setting to 282 seconds (intermediate, 84% reduction) and 230 seconds (minimal, 87% reduction). The largest speed-up was observed with AF3, which dropped from 1441 seconds in the maximum setting to 33 seconds (intermediate, 97% reduction) and 29 seconds (minimal, 98% reduction), primarily due to skipping AF3-based MSA generation using JackHMMER.

To assess whether these runtime gains compromised model quality, we compared complex confidence scores based on pAE interaction (ipAE) and structural agreement (DockQ) to the twelve experimental structures across settings and structure predictors (Fig. **2**D). Importantly, both the intermediate and minimal settings maintained ipAE and DockQ scores similar to those of the maximum setting. These results demonstrated that our streamlined settings enable fast and accurate scoring of de novo designed binders seemingly without compromising model quality. Based on these findings, and the expectation that the minimal settings do not affect confidence scores for non-binders we adopted the minimal configuration for all subsequent analyses.

### Features that predict binding

We first aimed to identify which features from our scoring pipeline best discriminated between experimentally verified binders and non-binders across all data. The distributions of some features were significantly different between binders and non-binders, while other features had very similar distributions (Fig. S2, S3, S4, S5 and S6). We also observed that the template-based AF2 initial guess structures were distinct from those generated by ColabFold, Boltz-1 and AF3, which tended to produce more similar structures among themselves. This distinction was evident both in terms of RMSD between predicted structures and in structural metrics such as interface size, ΔSASA and interface nonpolar residue count (Fig. S7, S8).

We next quantified the ability of each feature to separate binders from non-binders using average precision (AP), which avoids the need to set arbitrary thresholds and is preferable to the area under the receiver operating characteristic curve (AUROC) given the strong class imbalance favouring non-binders (28). Using AP, we identified the best individual features and in addition examined the effects of feature interactions, by trying all pairwise products within each structure prediction models feature set (Fig. **3**A). For AF3, ColabFold and Boltz-1 the best individual features were ipSAE_min, ipSAE_max and LIS, all of which are scores based on the pAE matrix and corrected to only capture the high confidence binding interface. ipSAE is calculated similarly to ipTM, but only considers interchain residue pairs below a pAE cutoff (<10 by default). Similarly, LIS is calculated by applying a pAE cutoff to interchain contacts in the pAE matrix, and taking the average after inverting the filtered pAE values to a scale from 0-1. In contrast to the other structure predictors, for AF2 initial guess ΔSAP was identified as the best individual feature. We note that ipSAE and LIS were not calculated for AF2 initial guess, and that several of the binder-target complexes in our dataset were already preselected for ipAE and plDDT, potentially exhausting the predictive power of these metrics (Fig. S9 and S10). Overall, the best individual metric for *in vitro* binding prediction across all data is ipSAE_min from AF3.

**Figure 3.**
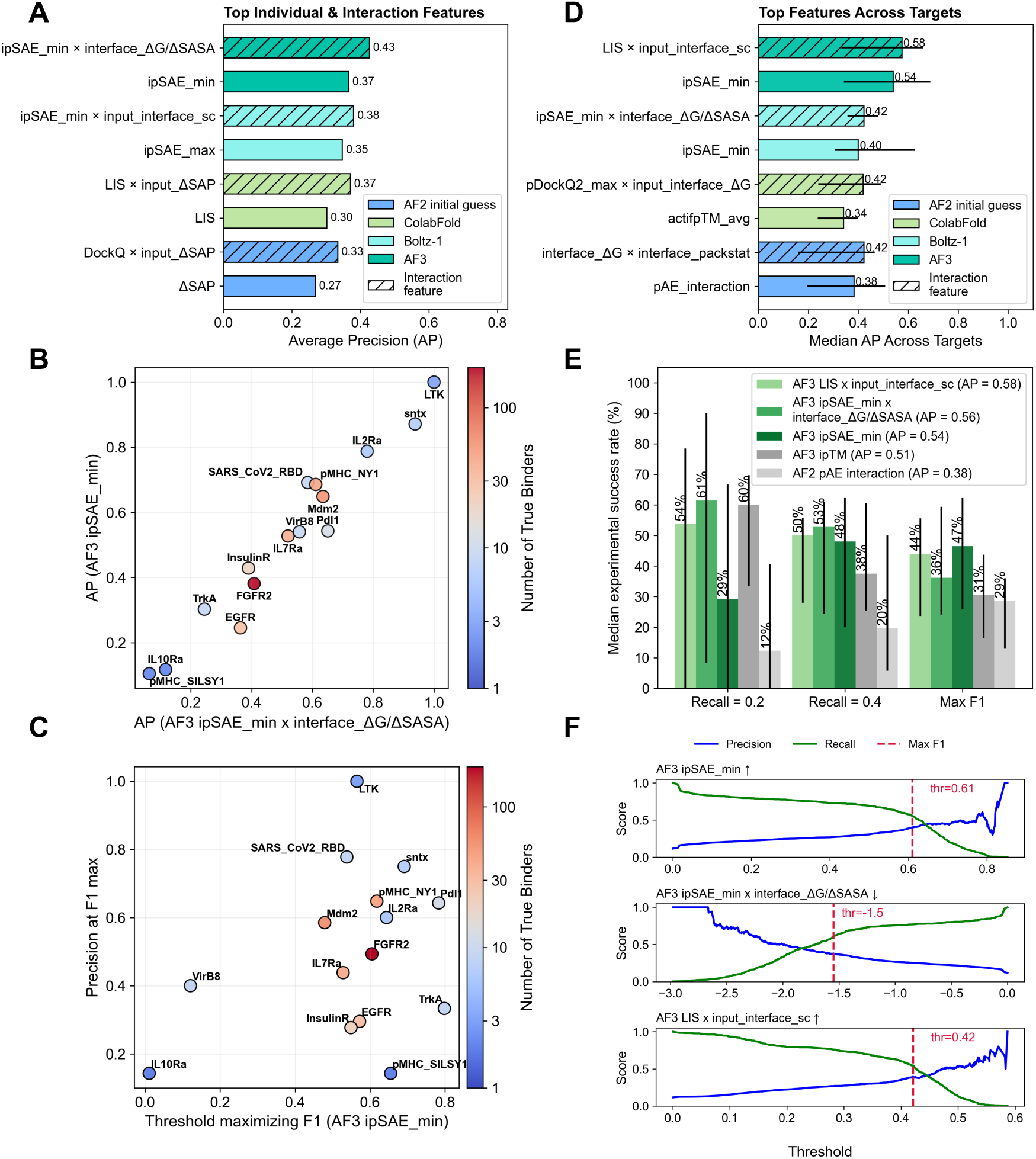
Single metrics that predict binding: **A)** Best individual (solid) and interaction (dashed) features for *in vitro* binder prediction across all data, as measured by average precision (AP). The best feature is shown for each structure prediction . **B)** Per target plot of AP of best individual and interaction feature from AF3 (AF3_ ipSAE_min x interface_ΔG/ΔSASA vs AF3_ipSAE_min). Each point represents a target, and is coloured by the number of positives binders. **C)** Per target plot of thresholds that maximize the F1 score for AF3 ipSAE_min and the corresponding precision at this threshold. **D)** Median AP of best individual (solid) and interaction (dashed) features for *in vitro* binder prediction across targets. Error bars represent inter quartile range across 15 targets. **E)** Median expected success rate across targets using recall 0.2, 0.6 and maximum F1 as threshold strategies for best individual and interaction features, with AF2 ipAE and AF3 ipTM as reference. Error bars represent inter quartile range across 15 targets. **F)** Precision and recall curves against different threshold for the three best overall features. Maximum F1 score is shown as red dashed line labeled with the corresponding threshold at this point. Arrows in title indicate whether the feature value is positively or negatively associated with binding probability.

After evaluating the predictive performance of individual features, we hypothesized that model performance could be improved by introducing interaction terms, defined as the product of two features. For features *f_i_* and *f_j_*, the interaction term is given by:

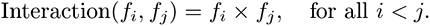

From the best individual features, the predictive power increases consistently across all structure prediction models when interactions between features are incorporated. The best combinations were ipSAE_min x interface_ΔG/ΔSASA, ipSAE_min x input_interface_shape_ complementarity, LIS x input_ΔSAP and DockQ x input_ΔSAP for the models AF3, ColabFold, Boltz-1 and AF2 initial guess (Fig. **3**A). Notably, for all structure predictors except the AF2 initial guess, the optimal feature combination included an interface-focused confidence score and a physicochemical interface descriptor, suggesting that these capture complementary information relevant to *in vitro* binding. Overall AF3 outperformed all other structure prediction tools for both the best individual and interaction feature indicating that AF3 potentially yields the most accurate confidence metrics and complex structures.

Next, we considered our targets independently as binding prediction and success rate in de novo binder design is known to be heavily target dependent, and because our dataset was imbalanced across targets, creating a bias towards targets with many designs. The AP of the best individual and interaction feature (AF3 ipSAE_min and AF3 ipSAE_min x interface_ΔG/ΔSASA), varied substantially across targets from 0.1 to 1. Targets with few true binders were found to be outliers, likely reflecting statistical variability (Fig. **3**B). We further examined how consistent AF3 ipSAE_min corresponded to precision across targets. Notably, the threshold of AF3 ipSAE_min that maximizes F1 (balancing precision and recall) fell within the range of 0.5 to 0.8 for most targets (with some outliers). However, the corresponding precision varied widely across targets, ranging from 0.1 to 1 (Fig. **3**C). We also observe that the top ranked features across targets varied quite substantially (Fig. S11). Due to the observed variability in feature performance across targets, we next aimed to identify features that perform well across most targets. We ranked features by their median AP across the 15 targets, which reduces the influence of outlier performance on individual targets. Once again, the top individual features from all structure predictors were confidence metrics based on a masked pAE matrix as the best individual feature, ranking ipSAE_min for AF3 and Boltz-1, actifpTM for ColabFold and pAE_interaction for AF2 initial guess as top features per structure prediction model. As before AF3 ipSAE_min stood out as best single metric, despite the observed variability in AP across targets. Aligning with the results observed across all data, the predictive power could be improved by including interaction terms. Again adding physicochemical interface descriptors, namely input_interface_shape_ complementarity (AF3), interface_ΔG/ΔSASA (Boltz-1), input_interface_ΔG (ColabFold) and interface_packstat (AF2 initial guess), to confidence based metrics consistently improved AP. The inter quartile range across targets was significant, indicating still a large difference in "predictability" of *in vitro* binding to the 15 targets in our dataset (Fig. **3**D). We next investigated whether any target specific properties correlate with the observed variability in prediction performance, as this could inform how tractable a design campaign is for a given target. We found that ΔSAP, a feature previously associated with target success rates due to its link to hydrophobicity, correlates with AP across targets when using AF2 ipAE. However, this correlation does not hold when using AF3 ipSAE_min, suggesting that the ipSAE score performs independently of ΔSAP (Fig. S12 A,B). Instead, AF3 ipSAE_min AP shows a positive correlation with the percentage of interface hydrogen bonds and a negative correlation with interface_ΔG/ΔSASA, suggesting higher predictability for targets with more energetically intense binding interfaces. However, these correlations are not statistically significant, and the observed trends are largely driven by a small number of targets (Fig. S12 C,D).

Based on the target-dependent differences, choosing the right threshold for filtering computational designs remains challenging, just as the corresponding experimental success rate is elusive. In a retrospective analysis for each of our targets, we tried three thresholding strategies for the best individual and interaction features and compared to the commonly used selection metrics AF2 ipAE and AF3 ipTM as baselines. Thresholds were determined via cross-validation: each target was held out once, and the threshold was selected based on performance across the remaining targets. We considered three practical thresholding strategies corresponding to: a recall of 0.2, a recall of 0.4, and the value that maximized the F1 score (balancing precision and recall). The individual feature AF3_ipSAE_min and the interaction features AF3_ipSAE_min x interface_ΔG/ΔSASA and AF3_LIS x input_interface_shape_complementarity consistently outperformed AF2’s initial guess metric ipAE across all thresholds. AF3 ipTM was also outperformed at recall 0.4 and max F1, but ranked second-best at recall 0.2, showing much lower interquartile range (IQR) than the top-performing interaction feature (AF3_ipSAE_min x interface_ΔG/ΔSASA) (Fig. **3**E).

To guide practical selections of thresholds for filtering de novo designs we compared precision and recall for the best three metrics across all data (Fig. **3**F). The best individual feature AF3_ipSAE_ min had a maximum F1 score at 0.61, whereas AF3_ipSAE_min x interface_ΔG/ΔSASA has max F1 at -1.5 (where lower score means higher probability of binding) and AF3_LIS x input_ interface_shape_complementarity had a max F1 at 0.42. To assess how the number of training targets influences threshold stability and performance, we performed a leave-one-target-out analysis on AF3 ipSAE_min, aggregating results across all possible combinations of training targets for each set size. Thresholds stabilized quickly, with precision and F1 improving consistently ( Fig. S13). All together these thresholds provide practical starting points for filtering *in silico* designs prior to experimental validation.

### Feature combination by greedy feature selection

After analysing which individual and interaction features best discriminated *in vitro* binders from non-binders, we aimed to identify whether using multiple features in a linear combination allows for improved selection of true binders. This was based on a twofold hypothesis: (i) agreement among confidence metrics from different structure prediction tools may carry meaningful information, and (ii) structural and physicochemical metrics may provide complementary, orthogonal insights to those confidence metrics. To this end, we established a greedy feature selection procedure using logistic regression in a Leave-One-Group-Out cross-validation approach, with each of the 15 targets as the holdout group. During greedy feature selection, features were selected in an inner cross-validation loop to maximize the median AP across the inner holdout targets, with early stopping of feature addition if AP did not increase more than 0.005 to prevent overfitting. The selected feature sets and corresponding logistic regression models were then evaluated on the outer holdout targets. We applied this procedure separately to individual features and to combined individual + interaction feature sets for each structure prediction tool. Interaction features were included to capture non-linear relationships that single-term models cannot represent. We also applied the procedure to a combined feature pool containing features from all structure prediction tools (Fig. **4**A).

**Figure 4.**
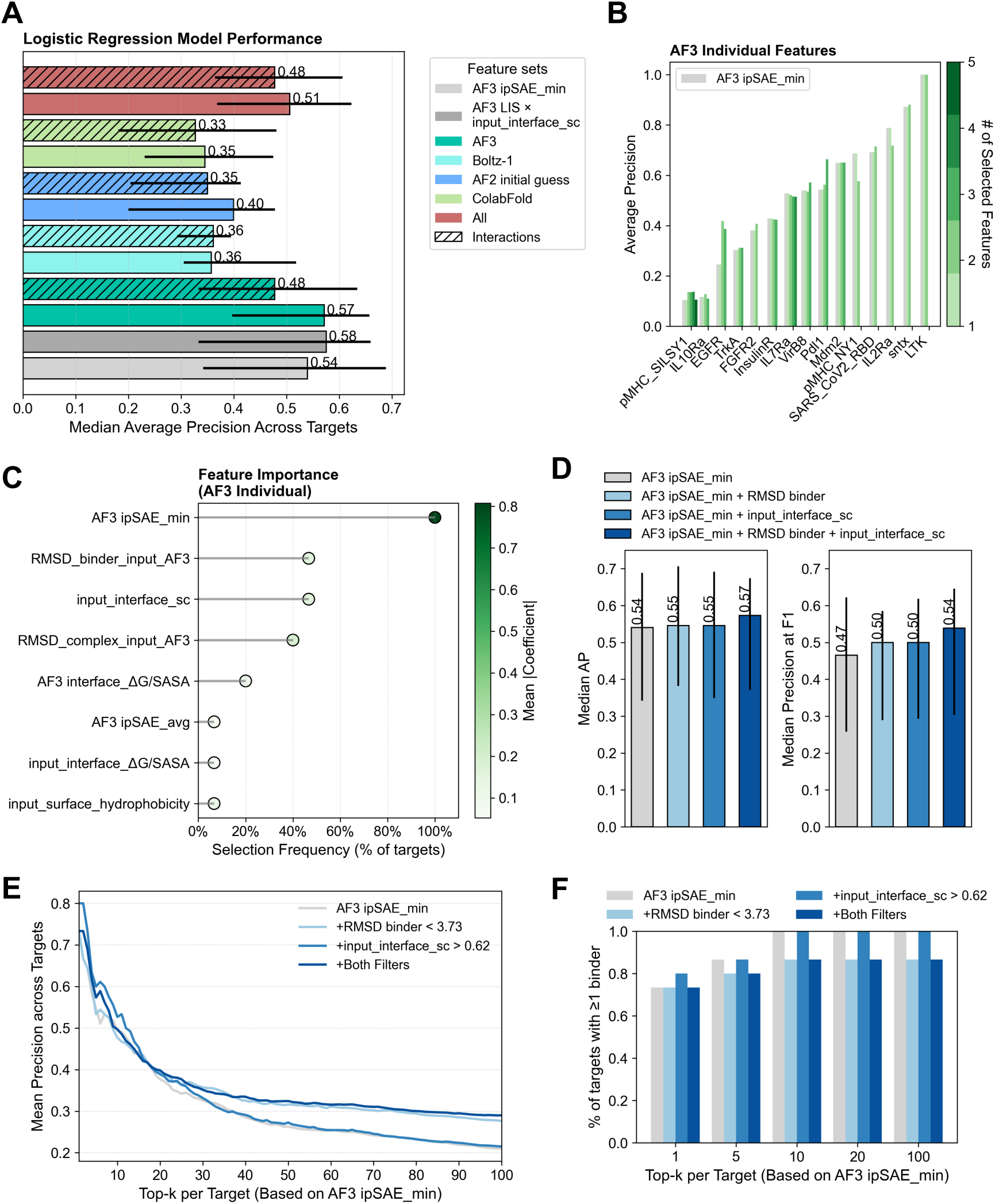
Greedy feature selection: **A)** Performance of logistic regression models using greedy feature selection and nested cross-validation based on the median average precision (AP) across holdout targets in the outer loop. Coloring corresponds to which feature set was used; "All" denotes the combination of features from all structure prediction tools, and hashed bars represent whether interaction features were included. Grey bars indicate the best single individual and interaction feature as baselines. Errors bars show inter quartile range (IQR) across all 15 holdout targets. **B)** Progression of AP on holdout targets in the outer loop for the model trained with AF3 individual features. **C)** Feature importance from models using AF3 individual features. Line length indicates how frequently a given feature was selected during greedy feature selection in the inner loop; dot color represents the mean logistic regression coefficient, reflecting feature importance. **D)** Median AP and precision at the F1-max threshold when retraining a logistic regression classifier using the top 3 features from C). **E)** Mean precision across targets when selecting the top-k binders based on AF3_ipSAE_min, with or without applying a RMSD_binder filter (< 3.73) and/or input_interface_shape_complementarity filter (> 0.62). F) Percentage of targets for which at least one true binder was selected at different top-k values, with and without filtering.

This analysis allowed us to assess whether features from different structure prediction models were equally informative, or whether combining features across models provided complementary value. The approach produced (i) an interpretable model in which feature importance can be inferred directly from the regression coefficients, and (ii) a feature set that only expands when features generalize well across different targets. To confirm that these results were not biased by redundancy in the dataset, we evaluated sequence-based similarity for both targets and binders. Apart from the expected similarity between the two pMHC targets and an observed similarity within the pMHC NY1 binders, both target and binder datasets exhibited a high degree of sequence diversity (Fig. S1 A,B).

While none of the models outperformed the baseline LIS × input_interface_shape_ complementarity (median AP = 0.58, IQR: 0.33–0.66) the model using individual AF3-derived features obtained a very comparable median AP (0.57) with reduced variance (IQR: 0.40–0.66), and it outperformed the best individual feature baseline AF3 ipSAE_min (Fig. **4**A). These two findings show that adding information from structural features can improve the prediction of the best confidence metrics, but the additional predictive power is easily exhausted. On average, only 2 to 5 features were added per model in the inner loop before improvement stagnated (Fig. S14 B), and when comparing performance in the outer loop to the best individual AF3 feature model, an increase in AP was observed only for a subset of targets (Fig. **4**B). Combining all features from every structure prediction model did not improve median AP, even though it was the top performer on a handful of targets (Fig. S14 A). These results suggest that incorporating features from other models seems to only introduce noise rather than add complementary information across targets. Similarly, adding interaction features did not lead to performance improvements, when predicting on unseen targets. Adding more features likely increases the models susceptibility to overfitting. Across all models less features got selected in the inner loop for models using interaction features (Fig. S14 B), indicating that most interaction terms do not generalize well across targets and likely provide redundant information. Consistent with these findings, switching to a more complex model such as XGBoost, which can capture nonlinear relationships, did not yield any performance improvement (Fig. S15 B). This reinforces the conclusion that only a small set of individual features generalize well across targets, and that the overall predictive signal is better captured by a simple linear model using these few informative features.

To investigate which features from the model using all AF3-derived individual features were most relevant and potentially contributed predictive power, we examined both the frequency with which features were selected in the inner loop and their average logistic regression coefficients (Fig. **4**C). Strikingly, AF3 ipSAE_min was selected in all cases, highlighting once again the robustness and predictive strength of this single feature. Beyond that, both the selection frequency and average coefficient values dropped sharply, suggesting that no other individual feature consistently improve prediction across targets. Notably, two features based on structural comparisons (C*α* RMSD) between the input and AF3 structure were frequently selected: one comparing the binder structures (after aligning the binder-target complex on the target) and one comparing the full complexes. Additionally, input_interface_shape_complementarity and interface_ΔG/ΔSASA were among the selected features supporting our earlier findings that these physicochemical interface descriptor can add complementary information to confidence based features.

Since the nested cross-validation produces different models for each hold-out target, we next aimed to build a single model using only the most frequently selected features and evaluate whether this could improve *in silico* filtering. We selected the three top features from the model trained on AF3 individual features: AF3 ipSAE_min, RMSD binder (input vs AF3), and input_interface_ shape_complementarity. Using these three features we retrained a logistic regression model as before, and assessed the median average precision and median precision at the threshold maximizing F1 across holdout targets (Fig. **4**D). Notably, adding both features to AF3 ipSAE_min had the highest increase in AP and precision and F1 max compared to just adding one of the features separately. This shows that adding a few additional features to AF3 ipSAE_min can in fact increase experimentally success rates.

To turn these findings into practical recommendations, we identified thresholds for the raw feature values that maximize the F1 score: RMSD_binder (< 3.73) and input_interface_shape_ complementarity (> 0.62). However, due to the observed high variability of optimal AF3_ipSAE_ min thresholds across targets, we adopted a different strategy for this feature. Instead of thresholding, we evaluated top-k ranked binders (k = 1 to 100, reflecting typical screening sizes) based on AF3_ipSAE_min, leveraging its strong ability to rank binders over non-binders. We performed this analysis with and without pre-filtering candidates using RMSD_binder and/or input_ interface_shape_complementarity (Fig. **4**E). We observe that for low k-values (∼1-20), adding the input_interface_shape_complementarity filter does indeed increase mean precision across targets as well as using both filters. For higher values of k using either the RMSD binder or both filters increased the mean precision. Importantly, AF3_ipSAE_min alone or combined with the input_ interface_shape_complementarity filter was sufficient to recover at least one binder for all 15 targets with as few as 10 candidates per target (Fig. **4**F). In contrast, applying RMSD_binder as a filter resulted in no binders being recovered for two targets across all k-values, suggesting that while this feature can enhance predictive power, it may also be overly restrictive in some cases.

Together, these findings show that only a small number of features are consistently informative for identifying true binders across targets. In particular, we demonstrate that combining structural metrics with AF3_ipSAE_min can further improve binder selection, and we outline how these features can be applied in practice through simple, interpretable filtering strategies.

## Discussion

De novo protein design is fundamentally changing the discovery of protein binders, enabling the efficient generation of binders against virtually any protein target from its structure alone. However, clear strategies for prioritising *in silico* designs for experimental testing are still lacking. In this study, we present a large-scale benchmark of structural, energetic, confidence, and sequence-based features for predicting binding success in de novo protein-binder campaigns. To this end we curated a dataset of 3,766 binder-target designs from 15 campaigns and re-predicted every complex using four structure prediction tools: the initial guess and ColabFold implementations of AlphaFold2 (AF2), AlphaFold3 (AF3) as well as its open-source implementation Boltz-1 (15–17, 27). For this we applied a streamlined prediction setup that reduced runtime by 87% while preserving interface-confidence accuracy. This enabled large-scale benchmarking and provides a practical, generalisable framework for evaluating confidence and structure-based metrics in protein binder design. Critically, it allowed us to identify metrics and combinations of metrics that correlate with binding success and improve the success rate over commonly used scores for binder selection.

AF3-derived metrics consistently show the highest predictive performance, even against the most comparable model, Boltz-1. This observation aligns with previous findings that AF3 outperforms other models in protein–protein interaction prediction tasks (17, 18). However, we note that the recently described Boltz-2 (18) and other models such as Chai-1 (19) may offer improved accuracy and warrant further evaluation.

We found that AF3 ipSAE_min was the best-performing predictor across all targets. The ipSAE metric (22) is a high-confidence-interface focused scoring scheme, and we specifically found that aggregating asymmetric predictions between chains using the minimum, rather than the average or maximum, yielded the most discriminative results. AF3 ipSAE_min provides a 1.4-fold increase in average precision (AP) compared to the AF2 initial guess pAE interaction (ipAE) score, which is commonly used to filter *in silico* designs of the state-of-the-art de novo design tool RFdiffusion (1). This improvement may stem from the fact that ipSAE focuses on high-confidence interfacial residue pairs and scales by interface size, leading to more consistent and target-agnostic performance compared to ipAE (22). We note that some data in this study had already been pre-selected using AF2 initial guess ipAE, potentially deflating the apparent predictive power of this metric. However, the ipAE score for Boltz-1, ColabFold and AF3 remained also suboptimal. The ipTM score, another commonly used scoring metric (e.g., in BindCraft (2)) had lower AP across targets compared to AF3 ipSAE_min, however it still performed well, especially at low recall thresholds. Our analysis further revealed that combining features through pairwise products can enhance predictive performance, by capturing interaction effects. Notably, the best performing interaction features combined interface focused confidence metric such as ipSAE or LIS (23) and a physicochemical interface descriptors such as the Rosetta based shape_complementary, ΔG/ΔSASA or ΔG. Consistent with this observation, a recent study focused on nanobody binding prediction also highlighted the importance of energy based metrics as predictive features (29). Together, these findings underscore the potential of integrating orthogonal feature types that are physically based, in addition to confidence scores to improve experimental success rates in de novo binder design campaigns.

Despite the improved predictive performance of individual and interaction metrics, we observed substantial variability across different targets. For AF3 ipSAE_min the precision ranged from 0.1 to 1.0 across targets at the maximum F1 threshold. We observed a higher AP of AF3_ipSAE_min for targets with high energy density in the binding interface as captured by higher H-bond percentage or more negative ΔG/ΔSASA ratio. Although the observed correlations with predictability were relatively weak, they may offer insights into the underlying factors that make some targets inherently more difficult to predict and design successful binders for. Notably, AF3 ipSAE_min AP did not correlate with interface hydrophobicity (ΔSAP), despite its known association with binder success (30). This correlation remained for AF2 ipAE, suggesting that AF3 ipSAE_min captures different aspects of interface quality and does not rely on hydrophobic burial.

We next evaluated whether combining multiple features in a linear fashion, rather than constructing interaction terms, could improve predictive performance across targets. We approached this using greedy feature selection with a logistic regression model and nested cross-validation, where each target was held out once and performance was evaluated using the median AP across holdout groups. This setup was designed to obtain an interpretable model that only selects features if they generalize well across targets. Notably, one model outperformed the best individual feature, but not the single best interaction feature in terms of AP across all targets, and most models selected only a small number of features. This suggests that only a few features consistently perform well across diverse targets and highlights the difficulty of building generalizable predictive models for scoring binders against novel targets. However, for the best-performing model using AF3-derived features, we found that incorporating interface shape_complementarity and RMSD values between the designed and AF3 re-predicted structures can improve experimental success rates. In particular, when selecting binders based on their top-k ranking by AF3_ipSAE_min, applying a shape complementarity filter to the designed input structures improved precision, especially for low k-values. This is particularly relevant as it provides a filtering step that can be applied prior to re-predicting the binder–target complex structure.

Based on our analysis, we recommend the following actionable filtering strategies. The first involves setting one of the following thresholds for filtering designs: AF3 ipSAE_min > 0.61; AF3 ipSAE_ min x interface_ΔG/ΔSASA < -1.5; or AF3 LIS x input_interface_shape_complementary > 0.42. The second strategy entails pre-filtering designs using input_interface_shape_complementary > 0.62 and RMSD_binder < 3.73, followed by selecting the top K designs based on AF3 ipSAE_ min.

We believe that further improvements can be made, if the limitations in the available data are addressed. For most targets only limited number of tested designs were available with only a handful of true binders reducing the model’s ability to generalize. Additionally, heterogeneous assay types and binding thresholds introduced label noise, and affinity data was only available for a small subset and thus not incorporated. More standardized, publicly available datasets linking binder structure to binding affinity are needed to improve predictive power and more fundamentally deepen our understanding of protein–protein interactions.

As a first step, our analysis demonstrates how to efficiently predict de novo binders and identifies which structure-prediction models perform best. Along with our actionable recommendations for candidate selection, the accompanying open-source dataset and pipeline provide a robust benchmark for the community. Together, these resources enable the systematic evaluation of structure-based binder scoring methods and will help accelerate the rapidly growing field of de novo protein design.

## Methods

### Collection of data

Datasets of miniprotein binders in complex with target structures with corresponding *in vitro* binding data were collected from different sources (Table 1) and compiled with the required information for our scoring pipeline (binder chain ID, target subchain ranges and MSA flags for each chain). All structures were gathered in PDB format with the binder as chain A and all target chains fused to chain B. For Bennett et al., and Cao et al., we sub-sampled a tenfold excess of non-binders compared to binders, as the full data sets exceeded what was feasible to run through our pipeline. Antibody binders from the Adaptyv bio dataset were excluded for this analysis to keep the focus on entirely de novo generated binders. Target interaction sites for true experimental binders were defined as antigen residues with any heavy atom within 4 Å of the binder. The frequency with which each residue formed part of an interaction site was calculated across binders and mapped onto the target structure for visualization in ChimeraX.

### Pipeline setup

Our pipeline uses input structures, i.e. the output binder-target complex structures from different de novo binder design tools (Figure **2**A). In addition, the pipeline requires an input CSV file, which describes the binder chain ID, target chain ID(s) and ranges as well as the MSA settings for each chain. This ensures that the input structures can have the binder and target in one chain each (which is required for AF2 initial guess) while the other models are aware of any sub-chains in the target. The output generated by the pipeline is a CSV file with one row per *binder_id*. All features in the output, both structural and confidence features are labeled with a prefix of the model from which they are calculated.

### Testing of different pipeline settings

To evaluate the impact of runtime-related parameters on prediction performance, we curated a benchmark dataset of 56 true binder–target complexes spanning all 15 targets plus three additional targets. Of these, 12 had experimentally determined structures that were used to assess model accuracy using pAE interaction (ipAE) and structural accuracy using DockQ (31). For each of these complexes, we computed ipAE and DockQ based on the highest-confidence predicted structure per model. The following PDBs were included in this evaluation: 7n3t, 7sh3, 7opb, 7n1j (30), 9bk5, 9bk6 (4), 8gjg, 8gji, 8t5e (13), 8wwc (32), 7xyq (33) and 9nnf (11).

The pipeline was tested under three configurations (maximum, intermediate, and minimal) which differed in MSA generation, number of recycles, and number of models per prediction (Fig. **2**B). In the maximum setting, paired MSAs were generated per binder–target pair using the default settings of each structure prediction tool (i.e. ColabFold’s MMseqs2 (34) for ColabFold and Boltz-1, JackHMMER (35) for AF3). In the intermediate and minimal settings, a single unpaired MSA was generated per target using ColabFold’s MMseqs2 implementation, reused across ColabFold, AF3, and Boltz-1, while binder sequences were predicted without MSAs. Additionally, in the minimal setting we reduced the number of models per prediction from five to three, and in ColabFold we cut the number of recycles from ten to three. All predictions were run on a single NVIDIA L40S GPU, and wall-clock runtimes were recorded for each configuration. The following repositories and versions were used: AF2 initial guess (https://github.com/nrbennet/dl_binder_design, v1.0.0), ColabFold with the AlphaFold-multimer (36) implementation (https://github.com/YoshitakaMo/localcolabfold, v1.5.5), Boltz-1x (https://github.com/jwohlwend/boltz, v1.0.0) and AF3 (https://github.com/google-deepmind/alphafold3, v3.0.1).

### Confidence scores

From each structure predictor the relevant confidence scores (from highest ranked model as well as average and standard deviation across sub models) were collected and the PDB file from the highest ranked model was extracted. Based on the confidence files (.npz files for Boltz-1 and .json files for ColabFold and AF3) the interaction prediction Score from Aligned Errors (ipSAE) (22) (using <10 pAE cutoff), Local Interaction Score (LIS) (average of inverted pAE scores in interchain contacts with <12 pAE) (23), pDockQ (37) and pDockQ2 (24) scores were calculated as described in (22). The ipSAE is calculated in a similar manner as ipTM, but instead of the interchain comparison employed in ipTM, it only included residue pairs below a certain pAE cutoff, and dynamically adjusts the *d*_0_ parameter according to the number of residues within the cutoff. The value of *d*_0_ increases with the square root of the number of interfacing residues. In turn, this means that a longer interface (with similar pAE values) results in a higher ipSAE score, which down-weights small but confident interaction surfaces, and reflects that very short interfaces are unlikely to result in binding.

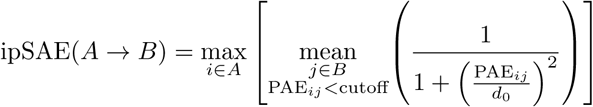

In addition to the ipSAE score versions previously reported (using different d0 corrections) (22), we also introduced some further modifications to ipSAE. Since the ipSAE score is asymmetrical, we stored both the max and min ipSAE score of any binder → target chain or target → binder chain value (denoted ipSAE_min and ipSAE_max). If the target had several subchains we took the average of the min and max values across both directions of binder → target comparisons, but only if there are interacting residues between the binder chain and a given target subchain. In the standard implementation (ipSAE_max), the asymmetric score is summarized by the max value, but we reasoned that the ’weakest link’ might be most informative for binding status. Furthermore, we also explored replacing max operator across chain A in the calculation of ipSAE, with an average or min, with the intuition of increasing the richness or stringency of the score respectively. We reported these scores as ipSAE_avg and ipSAE_min_in_calculation. We also calculated the ipAE from the pAE matrixes of Boltz1, ColabFold and AF3. This was calculated as an average of interchain pAE value from the binder chain to all other chains, following the implementation in (15).

### Structural features

For each pairwise combination of structure prediction models and towards the input PDB, we calculated C*α* RMSDs between models. This was done on the binder chain aligned on target chain(s) and on whole complex alignments referred to as RMSD_chA_aft_chB_align_model1_ model2 (or RMSD_binder_model1_model2) and RMSD_complex_model1_model2 respectively.

The Rosetta metrics were largely adapted from the BindCraft codebase (2). However, in addition to the metrics in the table below we calculated the probability of interaction using SpatialPPI V2 (38) on the input PDBs (i.e., spatialPPI_poi). We also calculated the CamSol intrinsic solubility scores of the binders based on input sequences (39). Further, we calculated the DockQ score between the input structure and each predicted structure. (31).

### Interface metric descriptions

#### Logistic regression classifier

All numeric features were normalized using z-score standardization with statistics from the current training split; class imbalance (∼12%binders) was handled with the inbuilt class_weight="balanced" which up-weighs the underrepresented true binders in the fit.

In the greedy forward-selection procedure, we left one target out at a time and used a Leave-One-Group-Out cross-validation over the remaining targets to choose features. Starting from an empty set, at each iteration we evaluated every candidate feature by training an l1-penalized logistic regression (with solver="liblinear") on the inner folds and measuring median average precision (AP). The feature whose inclusion yielded the largest AP gain was added, stopping once no remaining candidate improved AP by at least 0.005. After each feature was added, the model was also evaluated on the held-out target to track how performance based on AP evolved on the outer loop. Once selection finished, the final model was retrained on the inner data with C parameter optimization and evaluated on the outer set to report the final AP.

This greedy routine was run separately for each structure prediction model (AF2 initial guess, AF3, Boltz-1, and ColabFold), using their model-specific features together with the input structure features and general descriptors (binder/target length, spatialPPI, and CamSol score). For each case, models were trained either with all individual features or with all individual features plus the top 50 interaction terms (ranked by AP across targets) to reduce model complexity. In addition, two combined runs were performed, using either all individual features across models or all individual + interaction features across structure prediction models.

### XGBoost classifier

XGBoost models were trained using the same data-preparation and nested Leave-One-Group-Out cross-validation scheme as for the logistic-regression models. Briefly, all numeric features were median-imputed and z-score standardized within each training fold, and class imbalance (12% binders) was accounted for by setting scale_pos_weight = n_neg / n_pos in the XGBClassifier. In each outer fold one target was held out for final evaluation, while the remaining targets were used in an inner LOGO loop to tune hyperparameters (learning_rate, max_depth, subsample, colsample_bytree, min_child_weight) via grid search optimizing average precision (AP). Once the best parameter set was identified, the model was retrained on the full inner data and applied to the held-out target to compute AP.

## Data availability

The input PDB files, along with the outputs from AF2 initial guess, ColabFold, Boltz-1, and AF3 (including confidence files) and the compiled dataset containing all features generated in the analysis is available at doi:10.5281/zenodo.15722219.

## Code availability

Scripts for generating input files for the different structure prediction models, extracting all analysis metrics, and running the classification models is available at github.com/DigBioLab/de_novo_ binder_scoring.

## Acknowledgments

We would like to thank the HPC support team at the DTU Computing Center (DCC) for their valuable assistance in setting up the required infrastructure for this work. We further would like to thank Pia Haugaard Nord-Larsen and the administration team at DTU Bioengineering for their continuous support.

## Funding

T.P.J. and M.D.O. acknowledge support from the Alliance programme under the EuroTech Universities agreement. V.B. thanks the Novo Nordisk Foundation Center for Biosustainability (NNF20CC0035580) who funded this work. P.S. is a Royal Society University Research Fellow (grant no. URF\R1\201461) and acknowledges funding from UK Research and Innovation (UKRI) Engineering and Physical Sciences Research Council (grant no. EP/X024733/1).

## Author Contributions

M.D.O. and A.H. conceived the study. M.D.O. and A.H. curated the data, computed and evaluated all metrics, and established the logistic regression classifier. C.P.J., O.M., and V.B. assisted in setting up the training and validation of the classifiers. C.P.J. conducted the sequence similarity analysis. P.S. and T.P.J. supervised the study. M.D.O., A.H., and T.P.J. wrote the original draft of the manuscript. All authors reviewed and approved the final manuscript.

## Competing Interests

T.P.J. and O.M. are co-founders of AffinityAI. The remaining authors declare no competing interests.

---

**Fig. S1.**
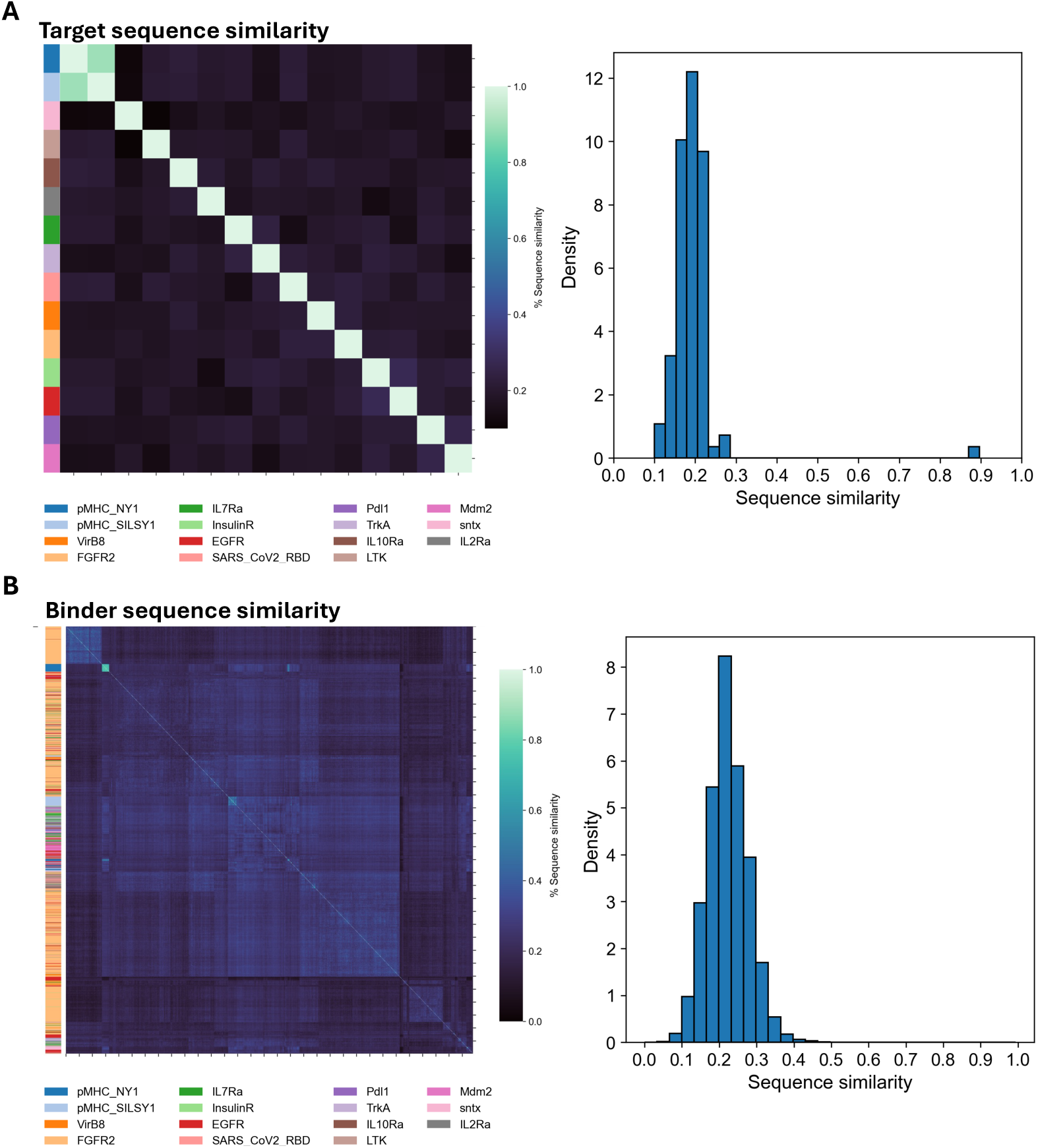
Sequence diversity across targets and binders: **A)** Clustered sequence similarity matrix based on normalized Levenshtein distances, with rows color-coded based on the target sequences. Histogram showing the normalized sequence similarity across targets. **B)** Clustered sequence similarity matrix based on normalized based on the binder sequences.Histogram showing the normalized sequence similarity across binder sequences.

**Fig. S2.**
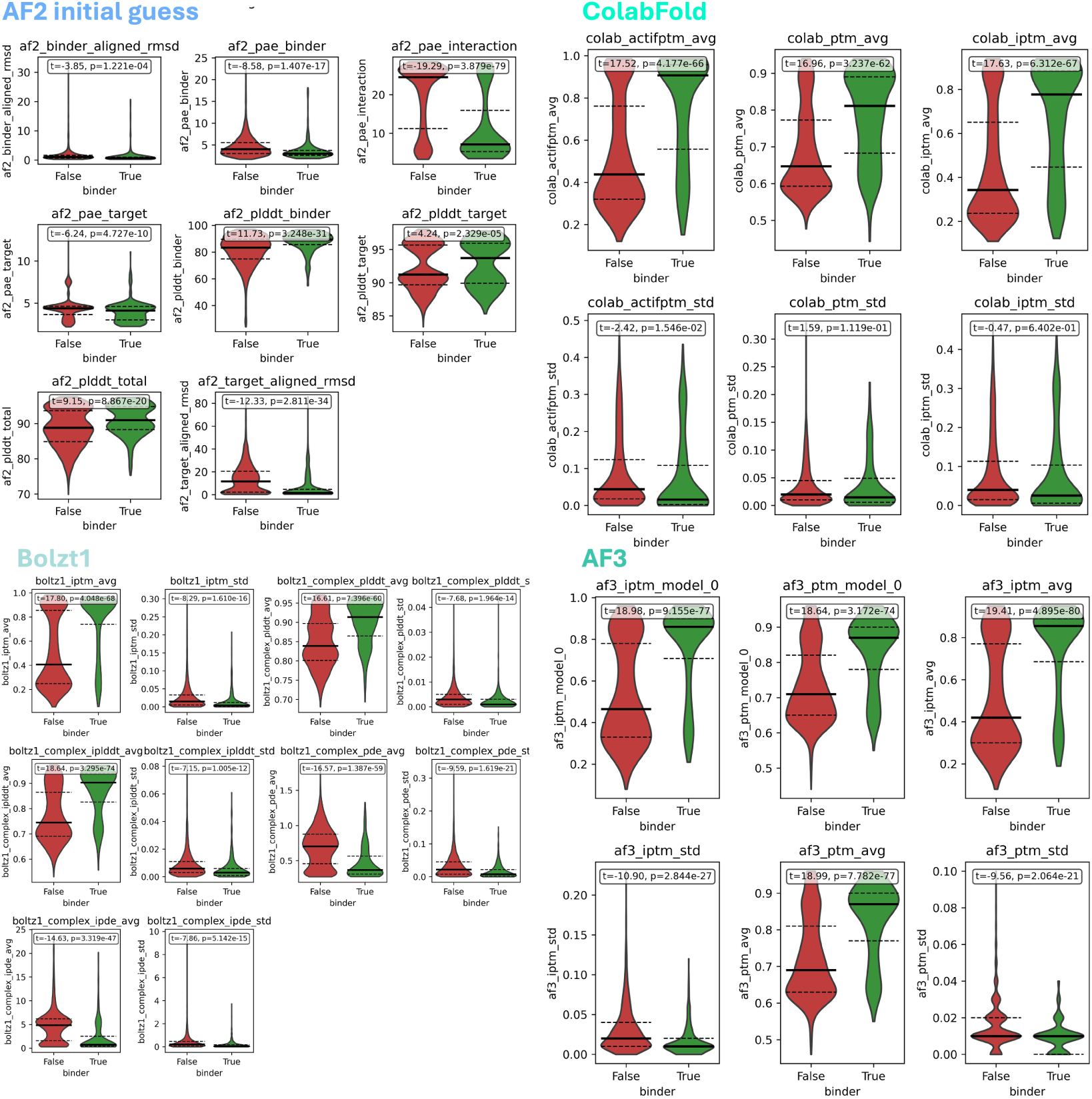
Violin plots confidence scores for binders vs non-binders: For each violin, the first quartile, median and third quartile are marked by black lines (median is thicker). P-value of an independent t-test between distributions is marked on each plot

**Fig. S3.**
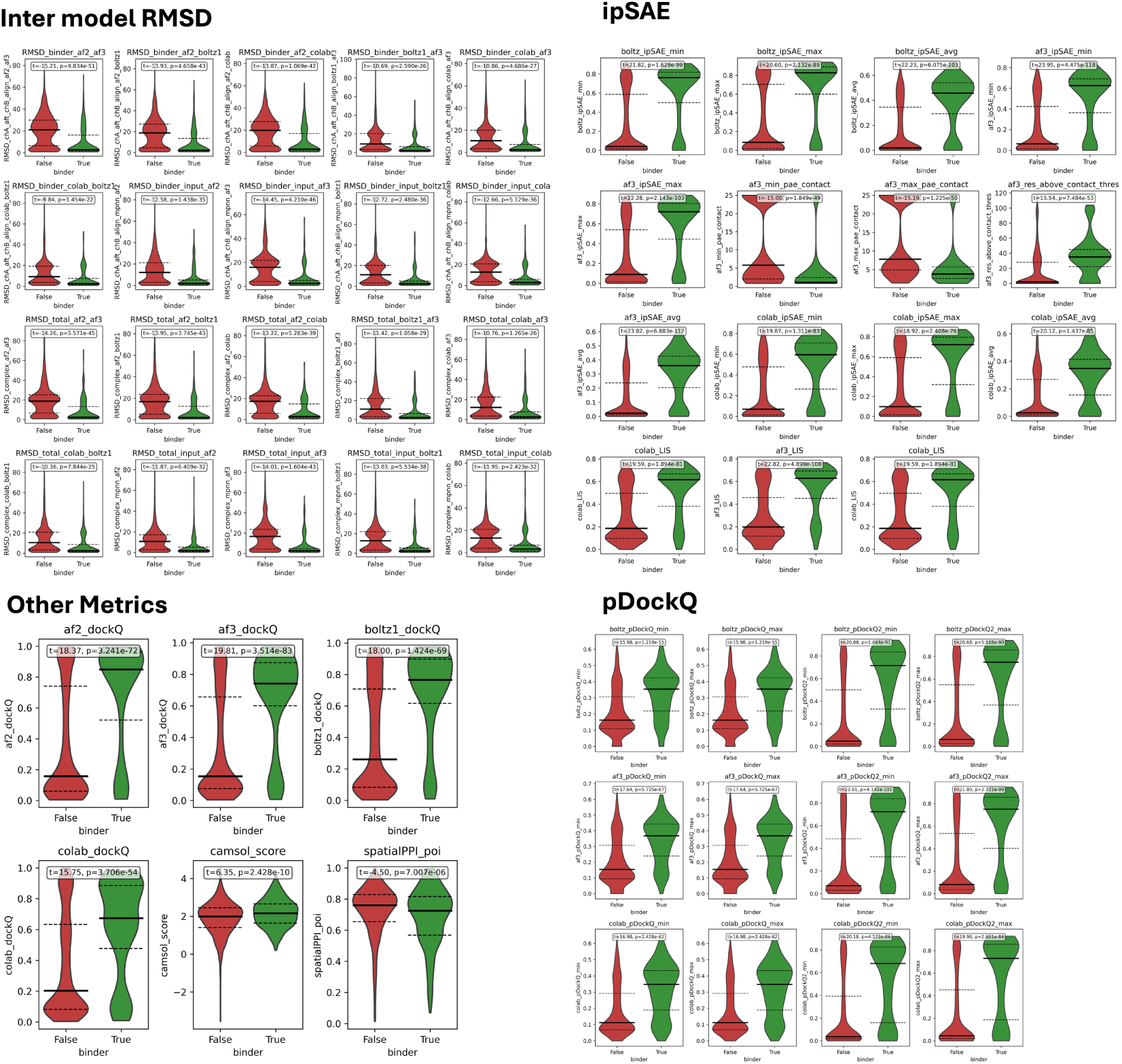
Violin plots of ipSAE, RMSD, pDockQ and other metrics for binders vs non-binders: For each violin, the first quartile, median and third quartile are marked by black lines (median is thicker). P-value of an independent t-test between distributions is marked on each plot.

**Fig. S4.**
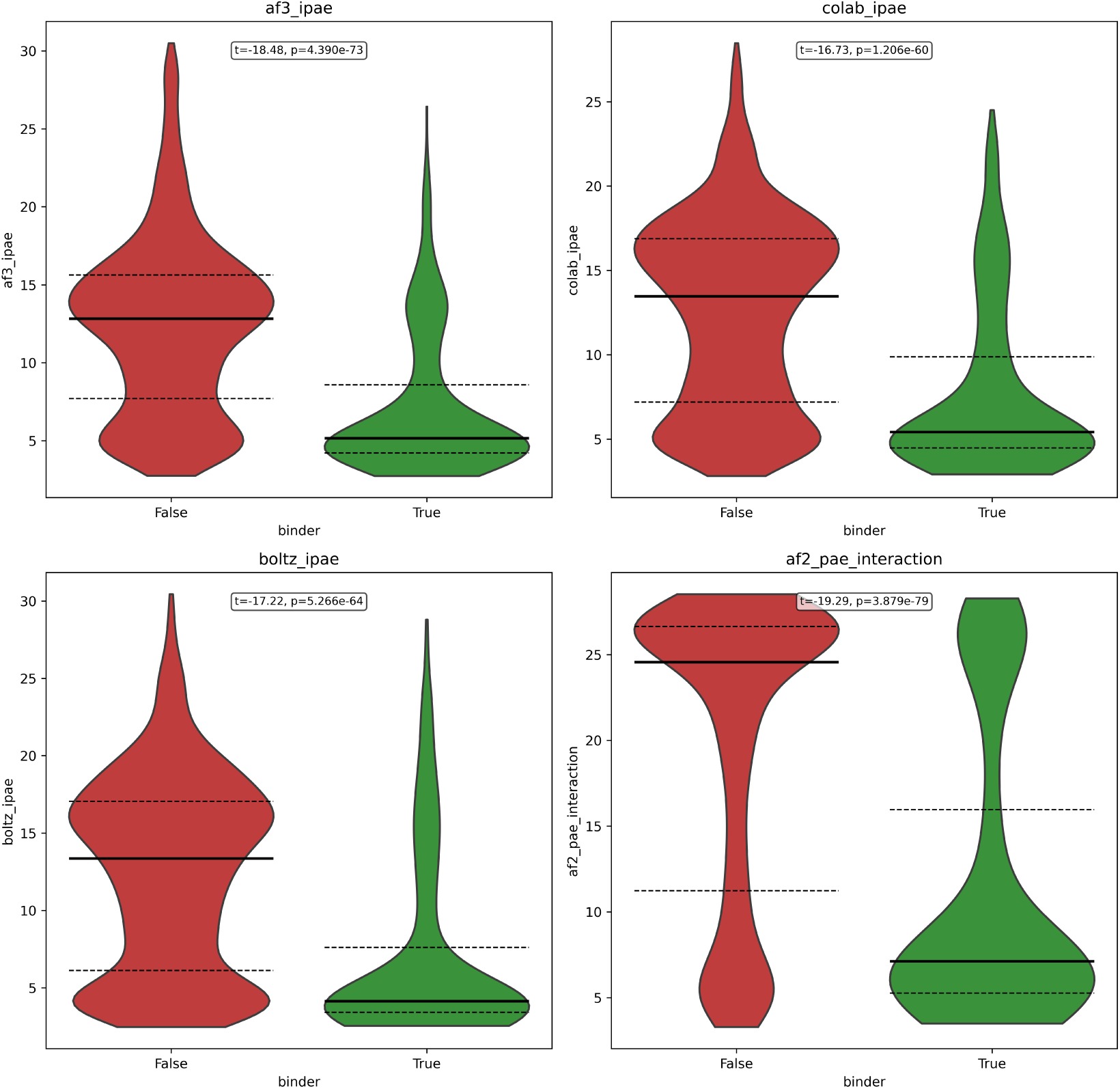
**Violin plots of ipAE**: For each violin, the first quartile, median and third quartile are marked by black lines (median is thicker). P-value of an independent t-test between distributions is marked on each plot.

**Fig. S5.**
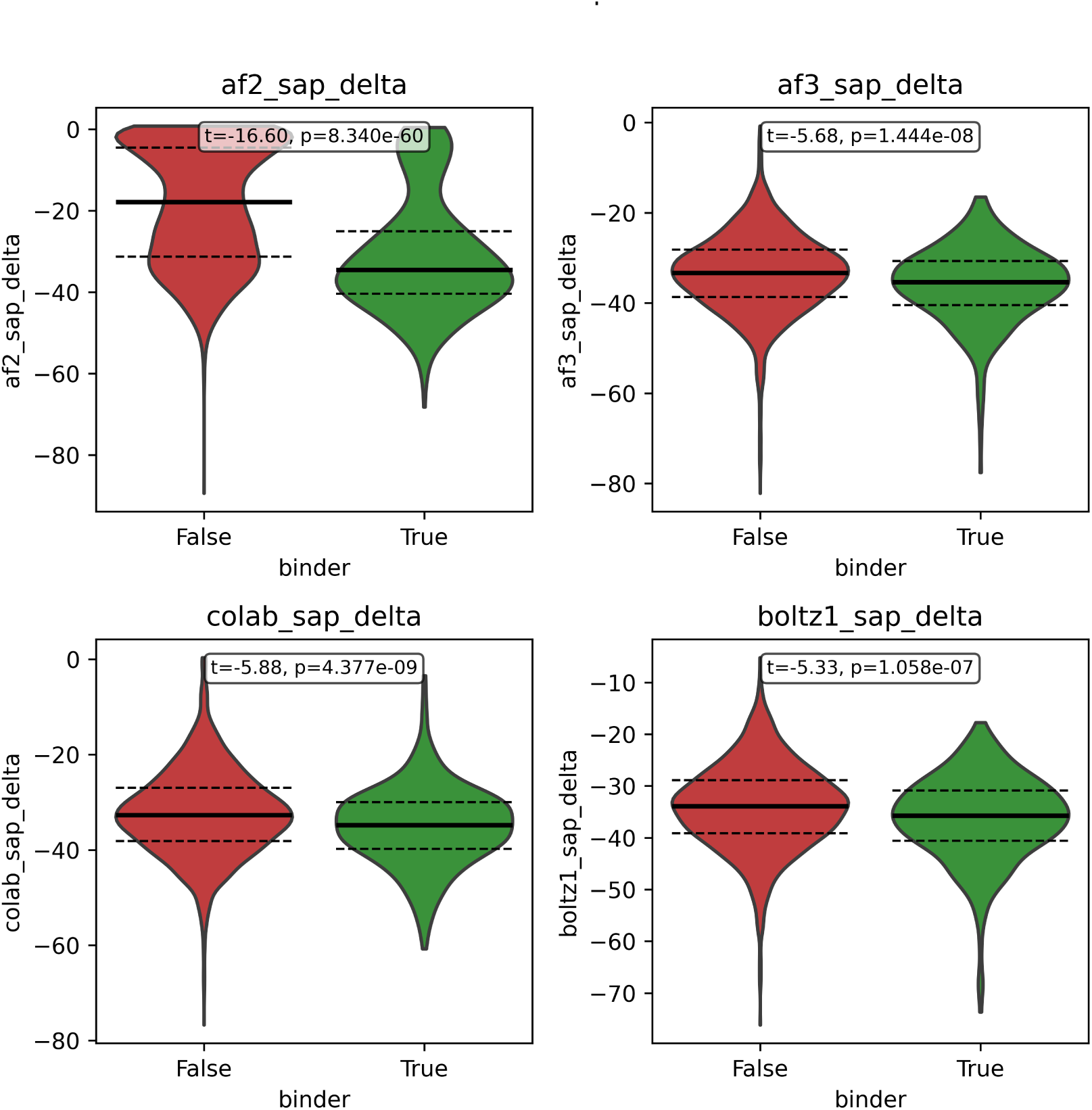
Violin plots of delta SAP for each model: For each violin, the first quartile, median and third quartile are marked by black lines (median is thicker). P-value of an independent t-test between distributions is marked on each plot.

**Fig. S6.**
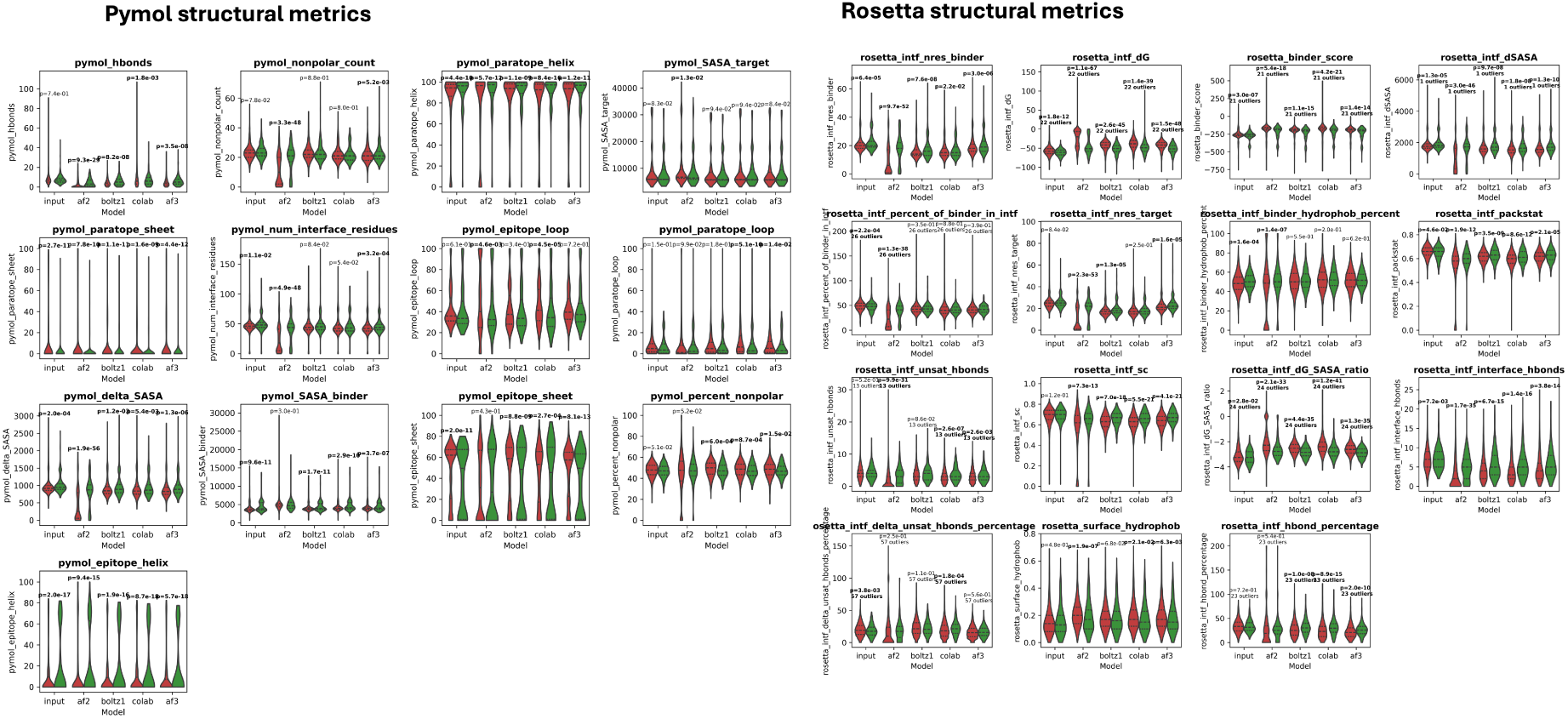
Violin plots structural metrics for binders vs non-binders: Structural metrics from rosetta and pymol are seen. For each violin, the first quartile, median and third quartile are marked by black lines (median is thicker). P-value of an independent t-test between distributions is marked on each plot.

**Fig. S7.**
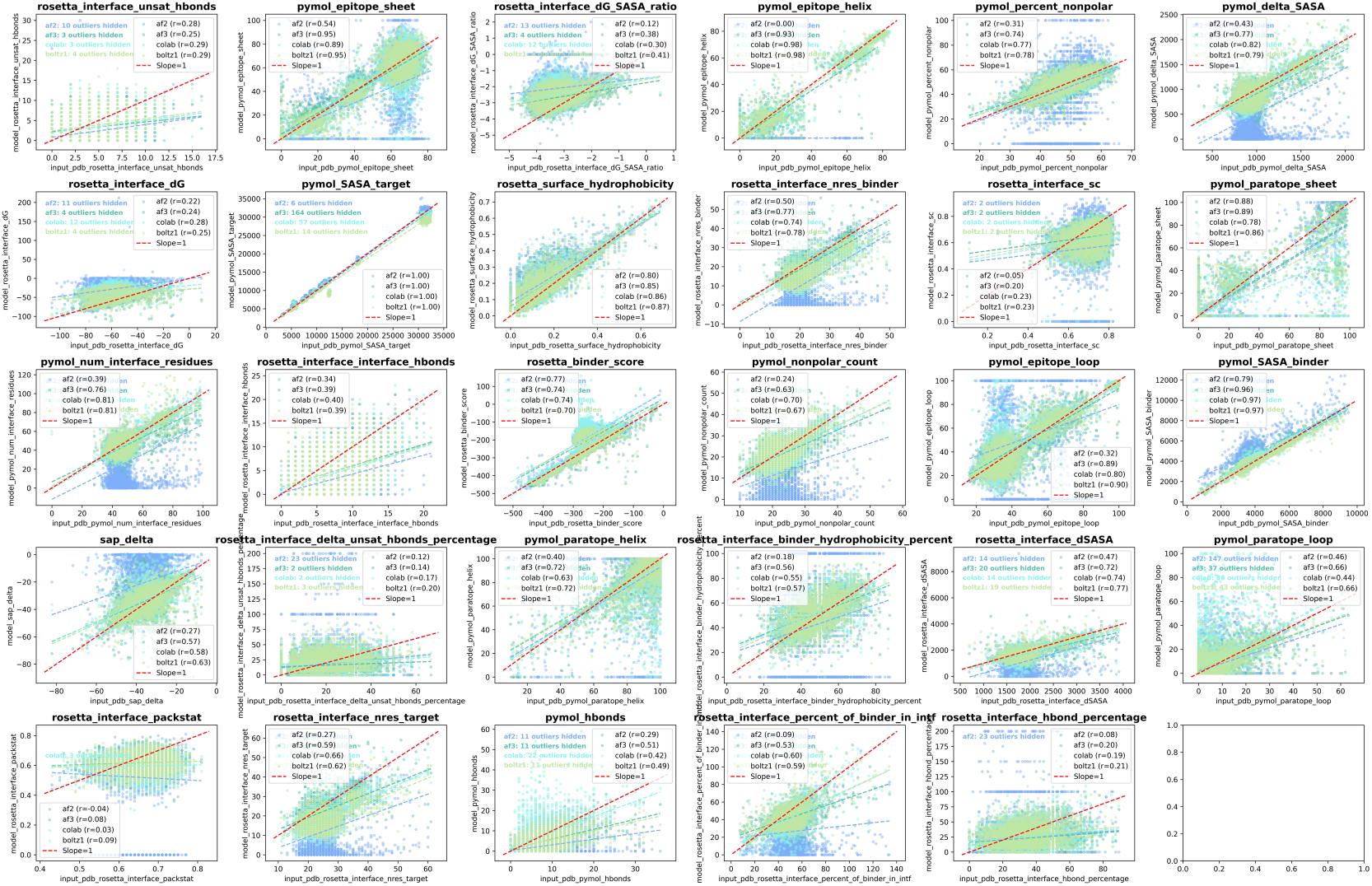
Comparison of structural metrics of input structure vs folding models structures: Scatter of all structural metrics from input PDB vs PDBs predicted by the folding models.

**Fig. S8.**
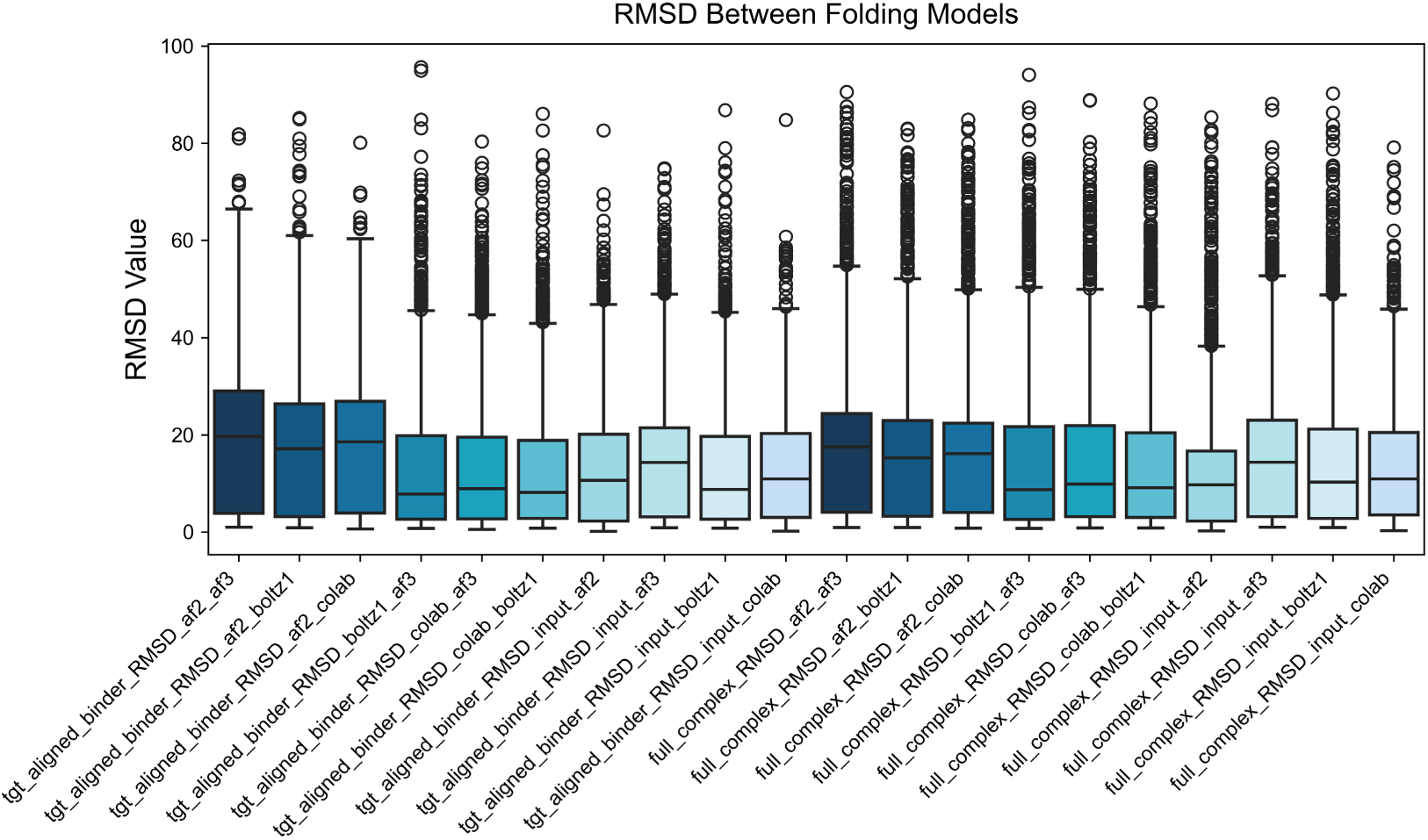
RMSD between structure prediction models: the C-alpha RMSD of binders after aligning the targets or whole complex alignments

**Fig. S9.**
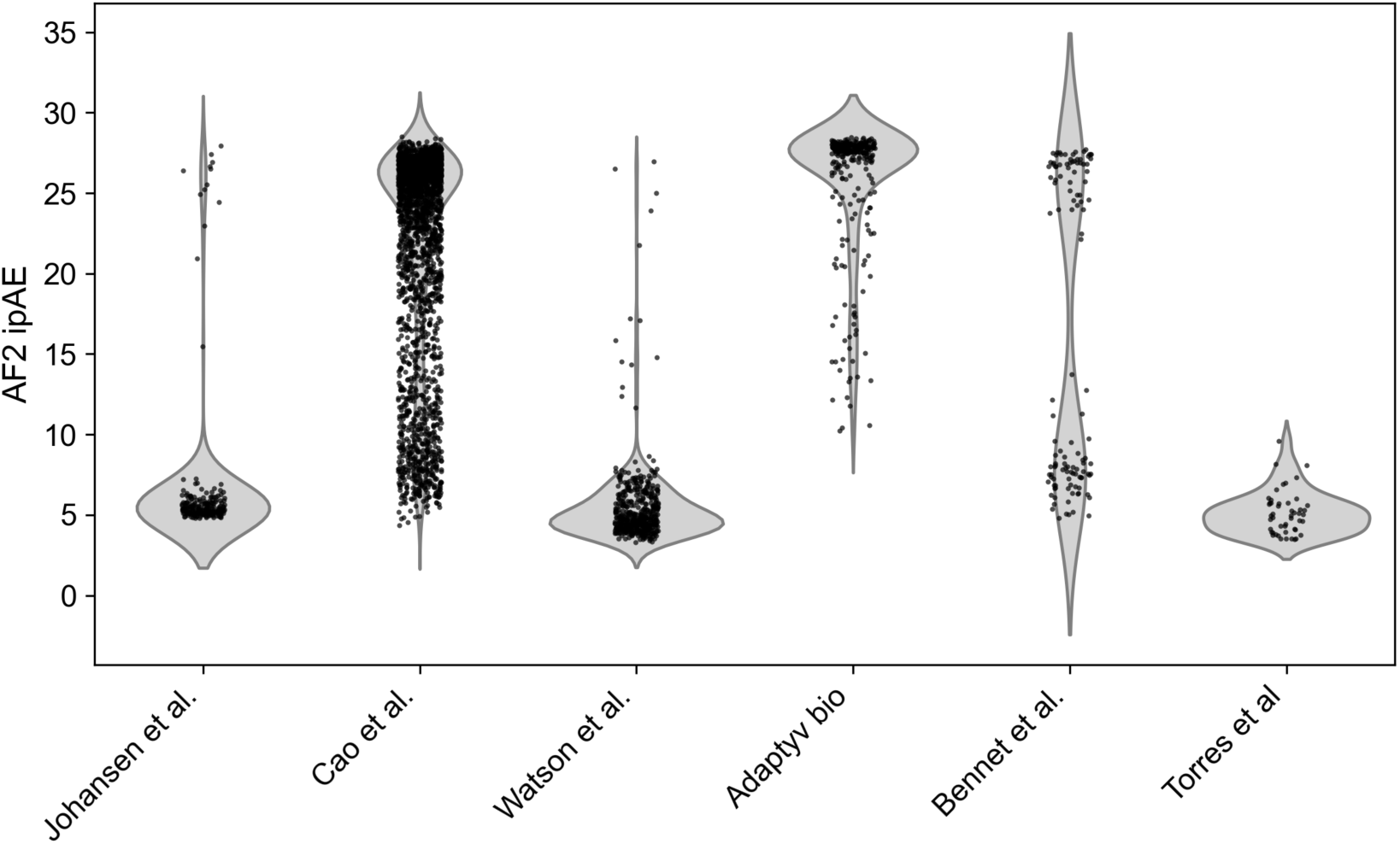
AF2 pae interaction across studies: Distribution of AF2 pae interaction (ipAE) across studies, highlighting that for several studies AF2 ipAE was used to filter *in silico* designs.

**Fig. S10.**
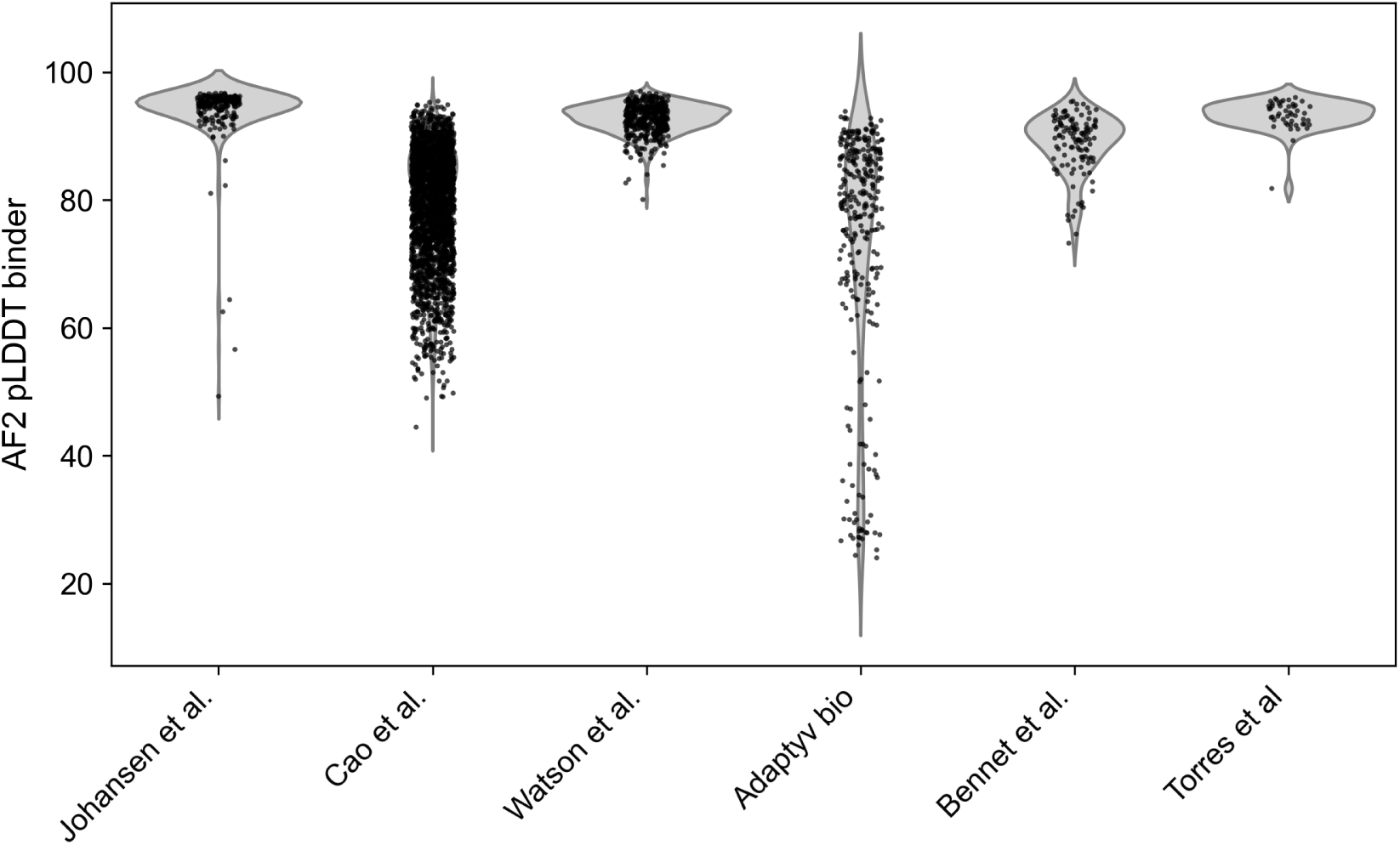
AF2 pLDDT binder across studies: Distribution of AF2 pLDDT binder across studies, highlighting that the distribution of AF2 pLDDT binder appears to skewed towards high values for some studies. Torres el. al directly states using pLDDT for filtering, while pLDDT is mentioned as a predictive metric in the retrospective analysis of Bennet et al., but is not used for selection (and Watson et al. follows the Bennet et al. recommendations of using only ipAE). Johansen el al. also only states using ipAE for filtering.

**Fig. S11.**
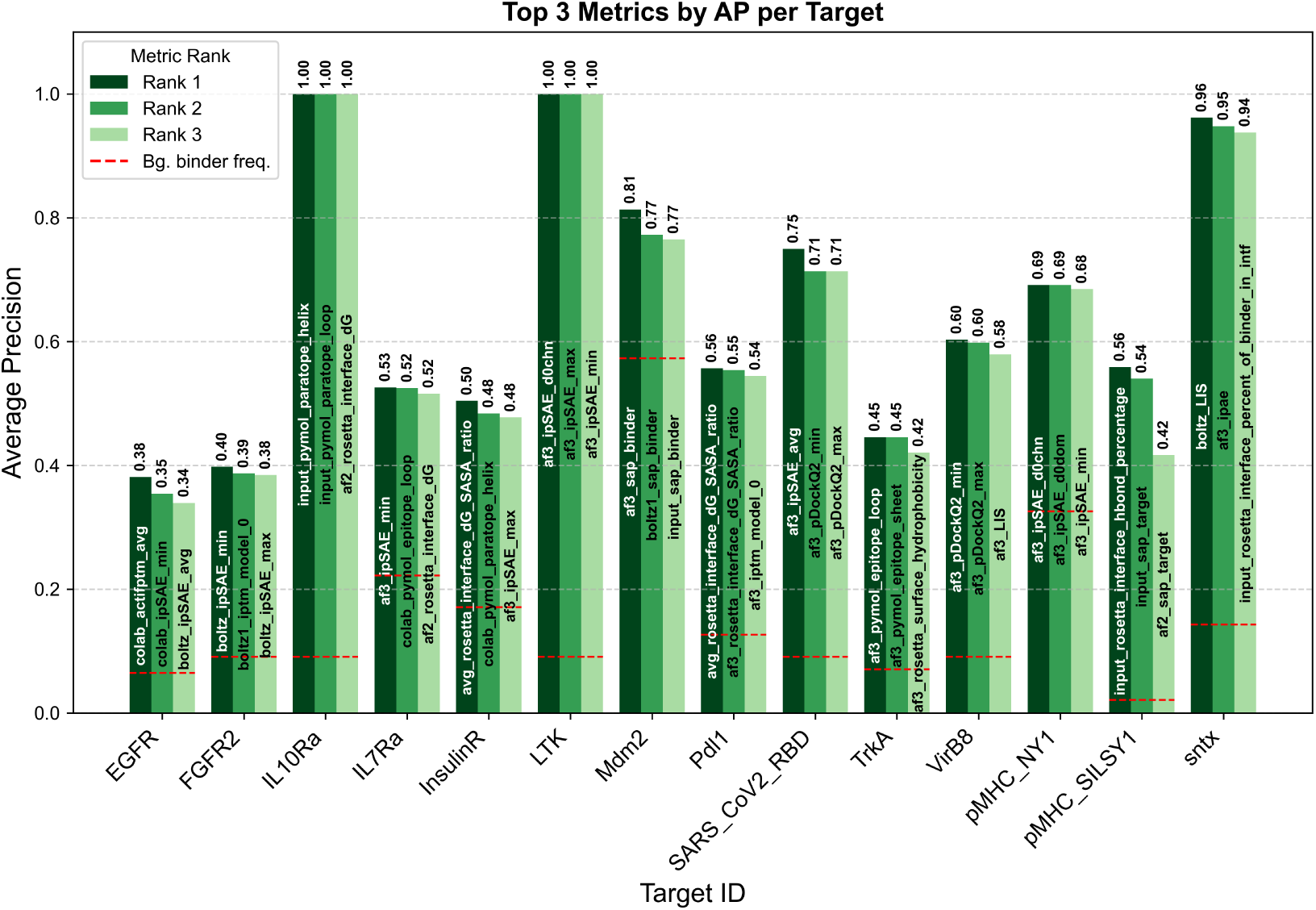
Top metrics per target: The best 3 metrics in terms of average precision is shown per target.

**Fig. S12.**
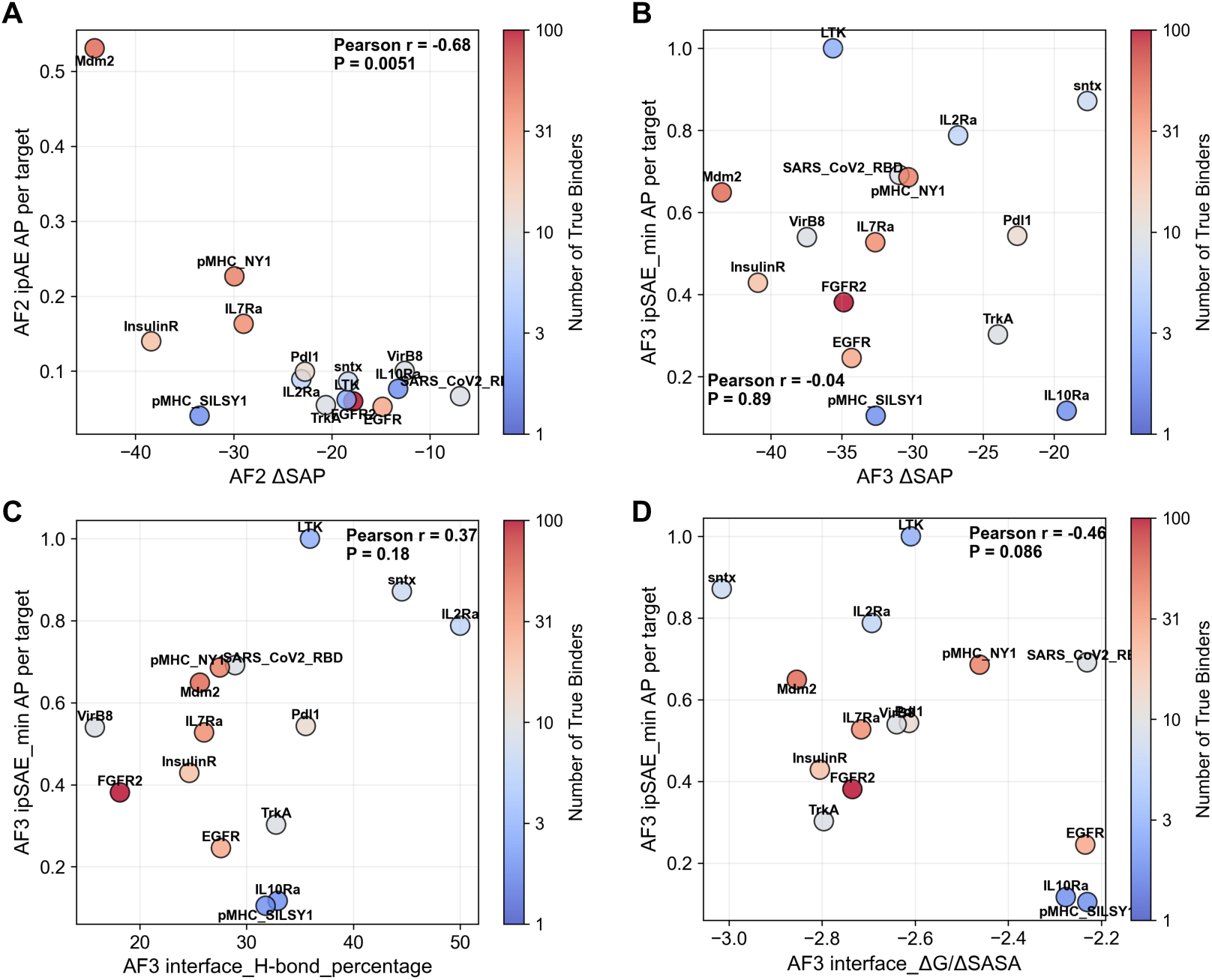
Assessment of predictability across targets: **A)** Correlation AF2 pAE*_i_nteraction*(*ipAE*)*averageprecision*(*AP* )*and*ΔSAP of AF2 initial guess predicted complexes across targets. For ΔSAP the mean across complexes for each target was taken. Coloring indicated number of true binder per target. Pearson R and two sided pval is shown. **B)** Correlation AF3 ipSAE_min AP and mean ΔSAP of AF3 predicted complexes across targets. **C)** Correlation between AF3 ipSAE_min AP and mean hydrogen bond percentage across targets. **D)** Correlation between AF3 ipSAE_min AP and mean ΔG/ΔSASA across targets.

**Fig. S13.**
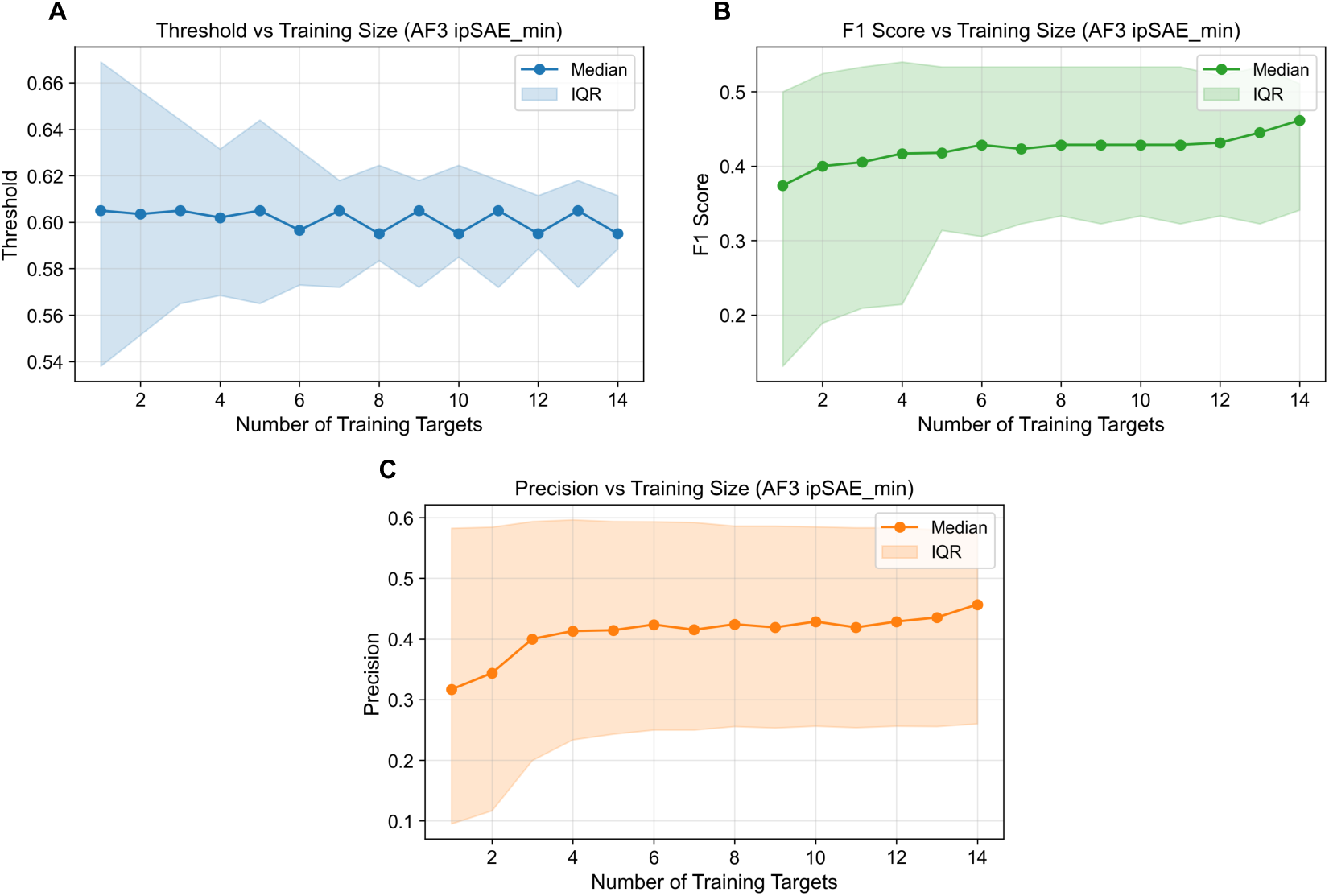
AF3 ipSAE_min threshold and performance stability: **A)** Threshold stability was evaluated using a leave-one-target-out approach on AF3 ipSAE_min, with thresholds for multi-target training sets defined as the median of per-target optimal thresholds (maximizing F1), and results aggregated across all possible combinations of training targets for each training set size. **B)** Corresponding F1-score and **C)** precision as a function of the number of training targets. IQR: interquartile range

**Fig. S14.**
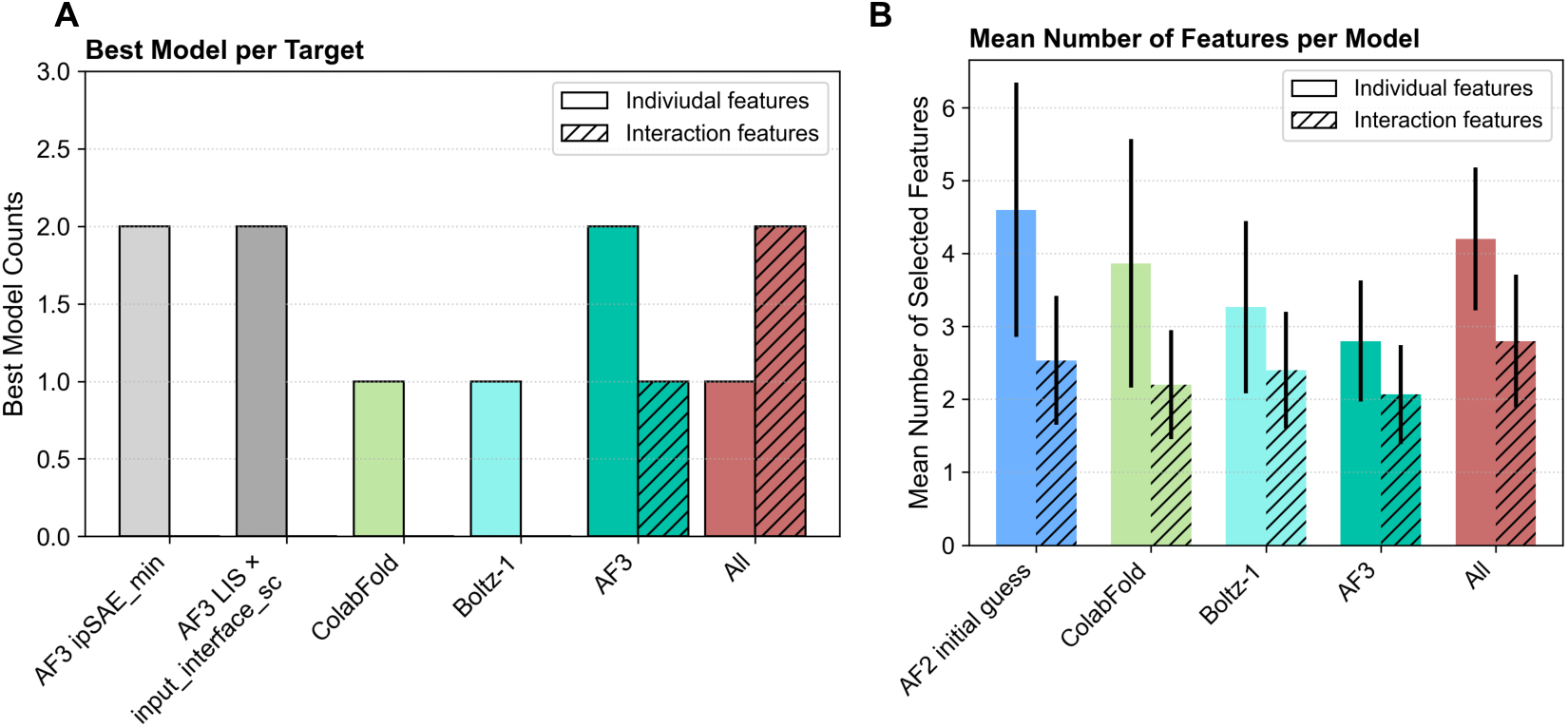
Logistic regression greedy feature selection evaluation: **A)** Per-target count of best performing model. **B)** Average number of features selected during greedy feature selection with a logistic regression using different feature sets, dashed bars indicate interaction features. Errors bars indicate standard deviation across all 15 holdout targets.

**Fig. S15.**
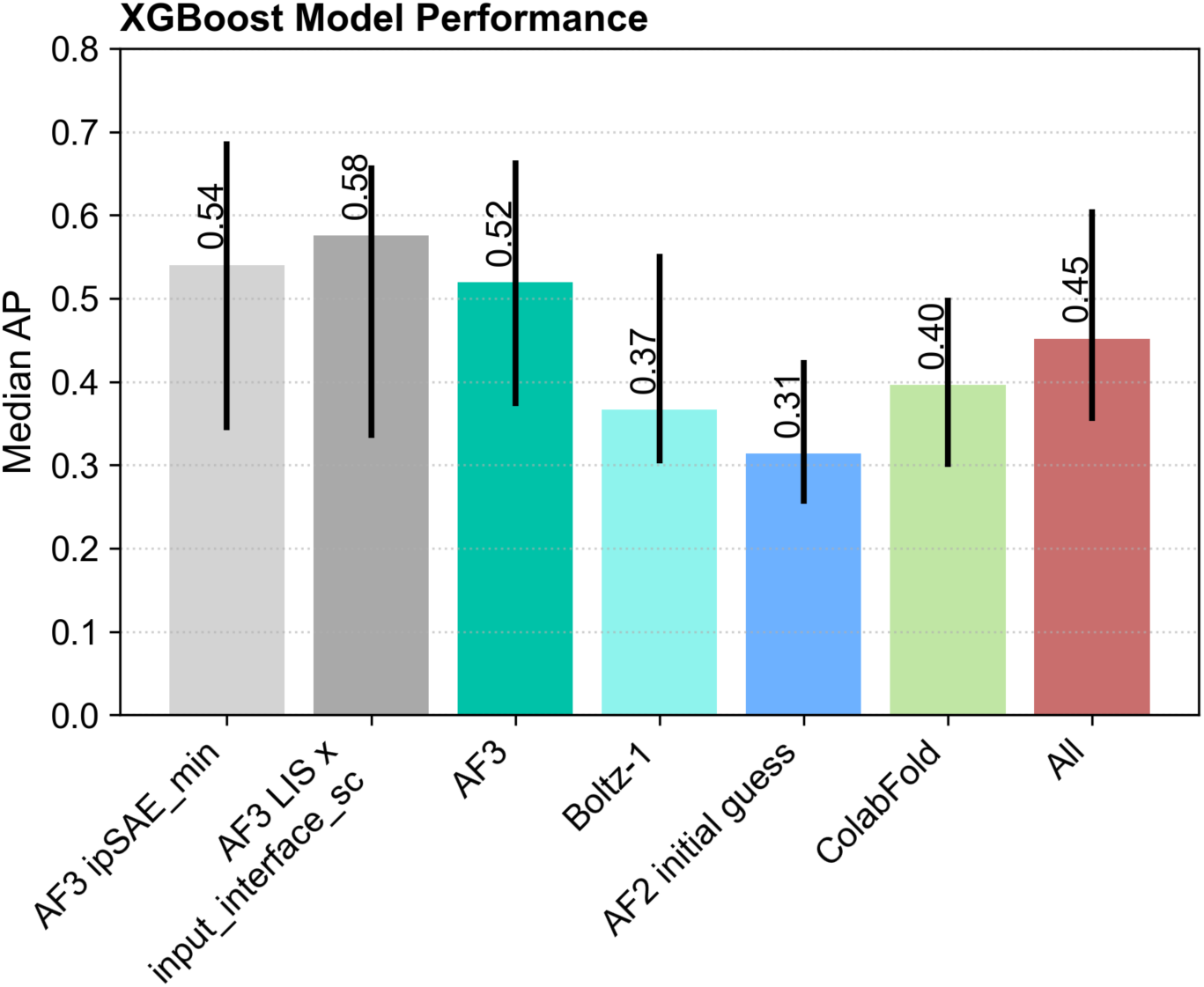
XGBoost model evaluation: Performance of XGBoost models with different feature set based on median average precision (AP) across holdout targets, error bars indicate interquartile range. No model performs better than the best baseline features AF3 ipSAE_min and LIS x input_interface_shape_complementarity.

## Notes

https://doi.org/10.5281/zenodo.15722219

https://github.com/DigBioLab/de_novo_binder_scoring

### Summary of Updates

Added github repo, made minor corrections (ipSAE formular, dG/dSASA defintion)

